# Cell-type specific developmental defects in *PTEN*-mutant cortical organoids converge on abnormal circuit activity

**DOI:** 10.1101/2022.11.15.516664

**Authors:** Martina Pigoni, Ana Uzquiano, Bruna Paulsen, Amanda Kedaigle, Sung Min Yang, Panagiotis Symvoulidis, Xian Adiconis, Silvia Velasco, Rafaela Sartore, Kwanho Kim, Ashley Tucewicz, Kalliopi Tsafou, Xin Jin, Lindy Barrett, Fei Chen, Ed Boyden, Aviv Regev, Joshua Z. Levin, Paola Arlotta

**Affiliations:** Department of Stem Cell and Regenerative Biology, Harvard University, Cambridge, MA 02138, USA; Stanley Center for Psychiatric Research, Broad Institute of MIT and Harvard, Cambridge, MA 02142, USA; Klarman Cell Observatory, Broad Institute of MIT and Harvard, Cambridge, MA 02142, USA; Decibel Therapeutics, Boston, MA 02215, USA; McGovern Institute for Brain Research, Massachusetts Institute of Technology (MIT), Cambridge, MA, USA; Murdoch Children’s Research Institute, The Royal Children’s Hospital, Parkville, Victoria, Australia; Broad Institute of MIT and Harvard, Cambridge, MA, USA; Society of Fellows, Harvard University, Cambridge, MA, USA; Department of Neuroscience, The Scripps Research Institute, La Jolla, CA, USA; MIT Center for Neurobiological Engineering, Massachusetts Institute of Technology (MIT), Cambridge, Massachusetts 02139, USA; Harvard-MIT Health Sciences & Technology Program (HST), Harvard Medical School, Boston, MA, USA; Koch Institute for Integrative Cancer Research, Massachusetts Institute of Technology (MIT), Cambridge, MA, USA; Howard Hughes Medical Institute, MIT, Cambridge, MA, USA; Department of Brain of Cognitive Sciences, Massachusetts Institute of Technology (MIT), Cambridge, MA, USA; Department of Media Arts and Sciences, Massachusetts Institute of Technology (MIT), Cambridge, MA, USA; Department of Biological Engineering, Massachusetts Institute of Technology (MIT), Cambridge, MA, USA; Department of Biology, Massachusetts Institute of Technology, Cambridge, MA, USA; Genentech, 1 DNA Way, South San Francisco, CA, USA

## Abstract

*De novo* heterozygous loss-of-function mutations in *PTEN* are strongly associated with Autism spectrum disorders (ASD); however, it is unclear how heterozygous mutations in this gene affects different cell types during human brain development, and how these effects vary across individuals. Here, we used human cortical organoids from different donors to identify cell-type-specific developmental events that are affected by heterozygous mutations in *PTEN*. We profiled individual organoids by single-cell RNA-seq, proteomics and spatial transcriptomics, and revealed abnormalities in developmental timing in human outer radial glia progenitors and deep layer cortical projection neurons, which varied with the donor genetic background. Calcium imaging in intact organoids showed that both accelerated and delayed neuronal development phenotypes resulted in similar abnormal activity of local circuits, irrespective of genetic background. The work reveals donor-dependent, cell-type specific developmental phenotypes of *PTEN* heterozygosity that later converge on disrupted neuronal activity.

## Main text

ASD is a childhood-onset neurodevelopmental disorder with cognitive, sensory, and motor deficits (Lord et al. 2020). ASD is characterized by strong clinical heterogeneity and a complex genetic component, with variants in hundreds of loci associated with increased risk of ASD (Geschwind and Levitt, Curr Opin Neurobiol, 2007; Grove et al., Nat Genet, 2019; Rosenberg et al., Arch Pediatr Adolesc Med, 2009; Ruzzo et al., Cell, 2019; Sanders et al., Nature, 2012; Satterstrom et al., Cell, 2020). Emerging evidence indicates that mutations in specific genes associated with ASD affect the developmental timing of specific cell types (Paulsen et al., Nature, 2022; Villa et al., Cell Rep, 2022, Lalli et al. 2020), and can vary in penetrance between genetic backgrounds (Paulsen et al., Nature, 2022). It remains unclear whether later in development there is convergence across genetic backgrounds on shared functional phenotypes.

To test the relationship between early and late phenotypes and their relationship to genetic backgrounds, we focused on heterozygous loss of function mutation in Phosphatase and Tensin Homolog (*PTEN*), which are strongly associated with ASD risk (Satterstrom et al., Cell, 2020). Moreover, prior work has associated *PTEN* with key cell-biological processes in neurodevelopment in mouse models, including roles in neural progenitor proliferation and neuronal structure, function and plasticity (reviewed in (Skelton et al., Mol Neuropsychiatry, 2020)). However, how *PTEN* mutation causes ASD risk remains largely unknown. While complete loss of function of *PTEN* has been associated with abnormal proliferation (Gregorian et al., J Neurosci, 2009; Groszer et al., Science, 2001; Li et al., Cell Stem Cell, 2017), the impact of heterozygous *PTEN* loss of function (as found in the context of ASD) in human brain development still needs to be elucidated.

Here, we used reproducible organoid models of the developing human cerebral cortex (Velasco et al., Nature, 2019) to investigate the cell-type- and dosage-specific phenotypes associated with heterozygous *PTEN* mutations across different donors (genetic backgrounds). We used CRISPR-Cas9 to generate induced pluripotent stem cell (iPSC) lines carrying a heterozygous protein-truncating frameshift mutation in the phosphatase domain, a region found mutated in patients (**Supplementary Fig. 1a**; (Orrico et al., Clin Genet, 2009)) (**Methods**), in two different genetic backgrounds (Mito210 and PGP1), and produced mutant and isogenic control organoids from these lines (**Supplementary Fig. 1b**). Protein quantification by Western Blot verified a reduction in PTEN protein levels in mutant iPS cells compared to isogenic control lines. Consistent with a reduction in PTEN function, the downstream effectors AKT and ERK (MAPK) showed increased phosphorylation (**Supplementary Fig. 1c**). Immunohistochemistry validated proper initiation of neural development in both *PTEN* heterozygous and isogenic control organoids (**Supplementary Fig. 1d, e**).

Because *PTEN* and its downstream effectors *AKT* and *MAPK1* are ubiquitously expressed across all cortical cell types during control organoid development by single-cell RNA-seq (scRNA-seq) (**Supplementary Fig. 1f**) (Uzquiano et al., Cell, 2022), we investigated the celltype-specific phenotypes caused by heterozygous mutation in *PTEN*, using scRNA-seq of individual organoids at one month in culture (**Fig. 1a-b** and **Supplementary Fig. 1g-h**). Analysis of 71,106 single cells from one batch each of Mito210 and PGP1 control and *PTEN* mutant organoids at one month, showed that *PTEN* heterozygous organoids had no significant difference in the proportion of any cortical cell type compared to control, in either genetic background (n = 3 single organoids per batch, FDR>0.05, logistic mixed model; **Supplementary Fig. 2a and b**). Thus, at early stages of development loss of one copy of *PTEN* does not cause differences in the proportion of any cortical cell type in human organoids.

**Figure 1.**
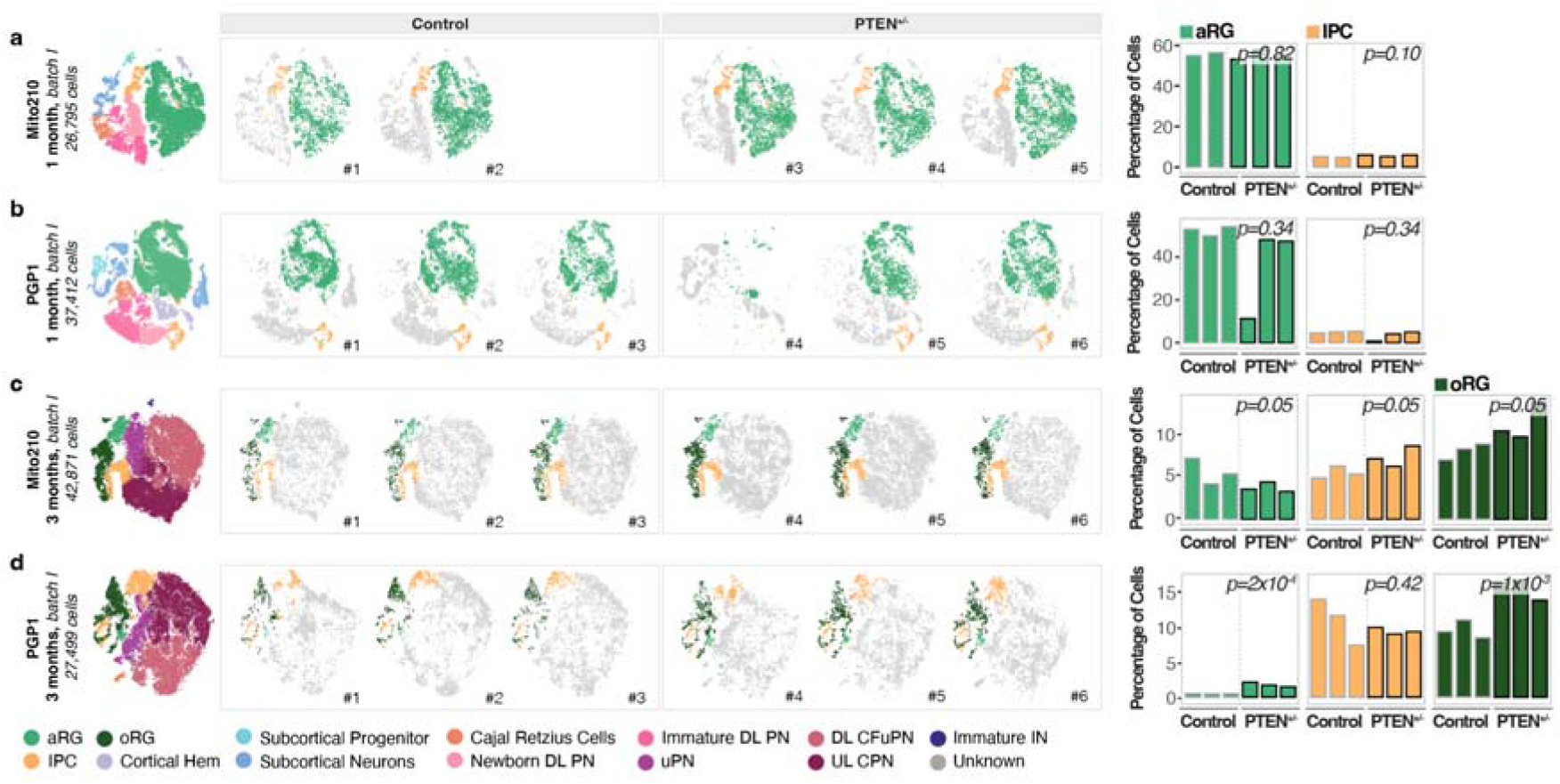
oRG are consistently affected in *PTEN* mutant organoids independently of genetic background. a-d) Left: t-SNE plots of scRNA-seq data, color-coded by cell type, from one month Mito210 control and *PTEN* heterozygous mutant organoids (Mito210 control n = 2, Mito210 heterozygous n = 3) (a), one month PGP1 control and *PTEN* heterozygous mutant organoids (n = 3 single organoids per genotype) (b), three months Mito210 control and *PTEN* heterozygous mutant organoids (n = 3 single organoids per genotype; batch I, see Supplementary Figure 2d for batch II) (c), and three months PGP1 control and *PTEN* heterozygous mutant organoids (n = 3 single organoids per genotype) (d). Middle: t-SNE plots for individual control and mutant organoids, with cell types of interest highlighted in color: apical radial glia (aRG, light green), intermediate progenitors cells (IPC, yellow) and outer radial glia (oRG, dark green). Right: bar charts showing the percentage of cells for the highlighted cell populations in each control and mutant organoid. FDRs for a difference in cell type proportions between control and mutant, based on logistic mixed models (see Methods) are shown. aRG, apical radial glia; DL, deep layer; UL, upper layer; PN, projection neurons; oRG, outer radial glia; IPC, intermediate progenitor cells; CPN, callosal projection neurons; CFuPN, corticofugal projection neurons; uPN, unspecified PN; IN, interneurons.

*PTEN* loss of function in early mouse development has been reported to affect the spatial organization of the mouse brain (Groszer et al., Science, 2001; Li et al., J Cell Biochem, 2003). We therefore investigated the effects of a heterozygous mutation in *PTEN* on spatial organization of organoid cell types at two months of culture, when this brain organoid model shows clear structural organization of progenitors into rosettes around the organoid periphery (Uzquiano et al., Cell, 2022), using spatial transcriptomics with Slide-seq on heterozygous and isogenic control Mito210 organoids. The spatial organization for most cell types was similar between conditions (**Supplementary Fig. 2c**), although outer radial glia (oRG), a progenitor cell type associated with human cortical expansion and evolution (DelValle-Anton and Borrell, Physiol Rev, 2022; Pinson and Huttner, Curr Opin Cell Biol, 2021), were more disorganized in mutant organoids, with a less well-defined ring-like arrangement of rosettes around the organoid periphery (**Supplementary Fig. 2c**).

This led us to wonder whether the composition of organoid cell types would be affected at later time points. We therefore profiled 102,216 additional cells with scRNA-seq from three separate differentiation batches at a later stage, 3 months in culture (n = 3 separate organoids per genotype per batch) (**Fig. 1c-d**, **Supplementary Fig. 1i-j** and **Supplementary Fig. 2d-e**), when the highest diversity of progenitor and projection neuron subtypes is found in these organoids (Velasco et al., 2019, Uzquiano et al., Cell, 2022). In both genetic backgrounds, heterozygous *PTEN* organoids showed a consistent significant increase in oRG (FDR<0.05, logistic mixed models; **Fig. 1c-d** and **Supplementary Fig. 2d**), whereas other progenitor types showed an increased proportion in heterozygous organoids in a background-dependent fashion: PGP1 *PTEN* mutant organoids displayed an increase in aRG (FDR=0.0002; **Fig. 1d**), and one batch of Mito210 *PTEN* mutant organoids showed an increase in IPCs (FDR=0.05 for batch I; FDR=0.02 for batch II; **Supplementary Fig. 2d**). Among projection neurons, one batch of Mito210 showed an increase in callosal projection neurons (CPN) (FDR=0.045; **Supplementary Fig. 2f-g**, Paulsen et al., Nature, 2022). However, the second Mito210 batch did not show a significant effect, while the PGP1 organoids showed a decrease in CPN (FDR=0.002; **Supplementary Fig. 2f-g**). No cortical cell type aside from oRG showed consistent differences in proportion in all three batches. Together, the data show that mutations in a single gene can have multiple cell-type-specific effects that can individually vary with genetic background.

Given the broad expression and function of *PTEN*, we wondered if later developmental events could also be affected by heterozygosity of this gene. We hypothesized that, despite the lack of consistent quantitative differences in proportions of the various neuronal populations in *PTEN* mutant organoids at one and three months, projection neurons may display alterations in their gene expression profiles or developmental speed across genotypes, as we and others have previously reported for other ASD-associated genes (Birtele et al., bioRxiv, 2022; Paulsen et al., Nature, 2022; Villa et al., Cell Rep, 2022, Lalli et al. 2020). Therefore, we compared the genes affected in *PTEN* heterozygous organoids to genes that vary in expression over organoid development, to relate transcriptional changes induced by *PTEN* mutation to developmental progression. We first calculated the differentially expressed genes (DEG) for each cell type between control and heterozygous mutant *PTEN* organoids at each time point (see Methods). We then used an existing high-resolution developmental atlas of human cortical organoids we previously obtained with the same differentiation protocol (Velasco et al., 2019, Uzquiano et al., Cell, 2022) to define gene changes associated with developmental time in each cell type. We calculated the genes that change over time in individual cell types across cortical organoid development, from all of the cell lines in that dataset, including the two used here (Mito210 and PGP1), as well as two others used previously (HUES66 and GM08330), depending on time point. From this analysis we obtained a list of genes that were up-regulated or down-regulated over time in specific cell types as organoids develop. We then compared the mutation-induced changes (i.e., DEGs between *PTEN* heterozygous vs control organoids in each cell type) at one month to the developmentally-regulated genes in the same subset from the atlas around this time point (i.e., DEGs that change over the period from day 23 through 1.5 months in control organoids in each cell subset). Similarly, the DEGs of *PTEN* mutant vs control organoids at three months in each cell type were compared to temporally-variable genes from the two through four months time period of the atlas for the corresponding types. To compare these lists, we used rank-rank hypergeometric overlap (RRHO2; **Fig. 2a-e** and **Supplementary Fig. 3a,c**; (Cahill et al., Sci Rep, 2018; Plaisier et al., Nucleic Acids Res, 2010)), which allowed us to determine whether the gene sets altered by the *PTEN* mutation indicated an acceleration or deceleration of development for each cell type.

**Figure 2.**
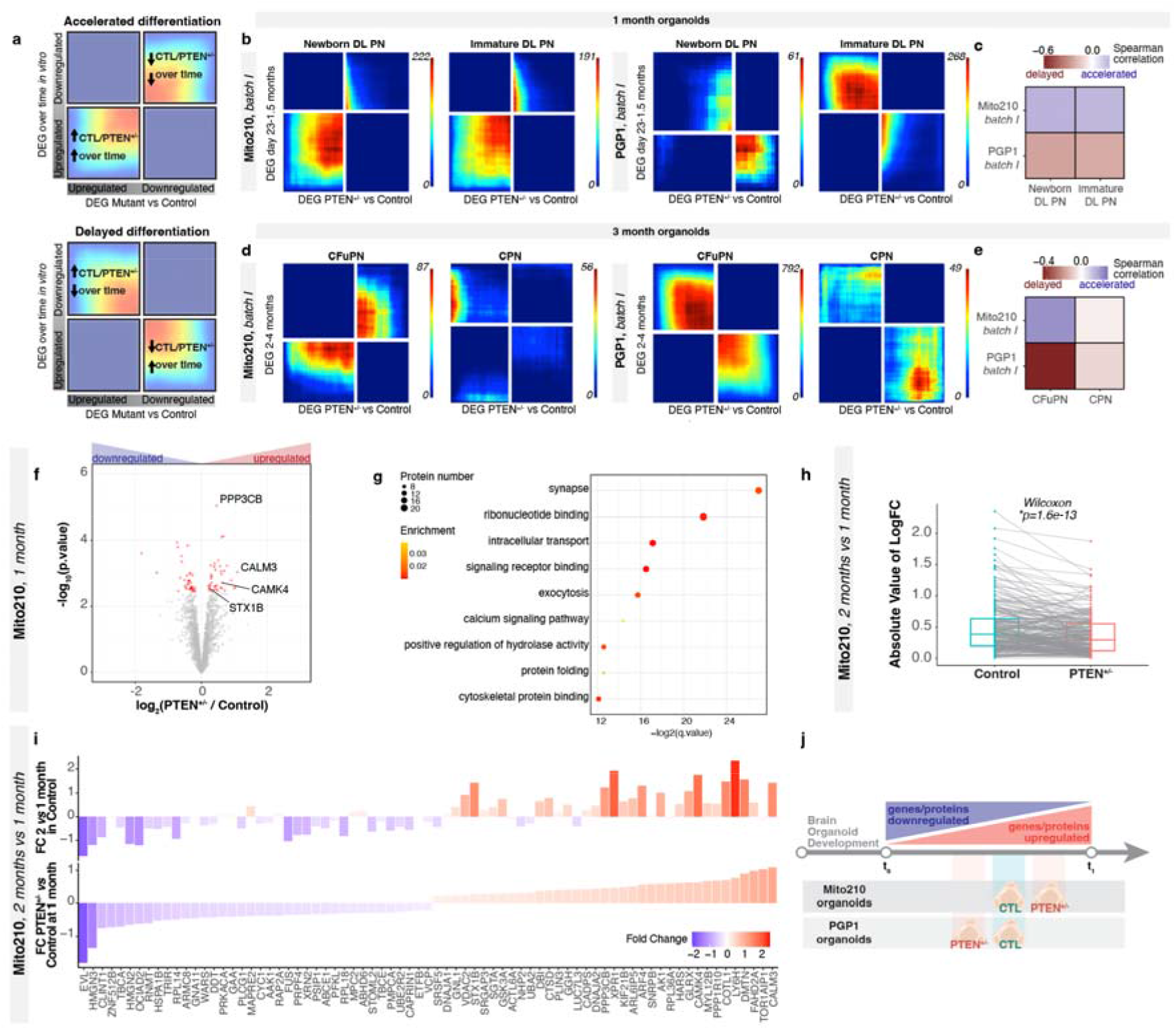
Deep layer neurons show asynchronous development across *PTEN* mutant organoids at multiple timepoints. a) Schematic explaining the rank-rank hypergeometric overlap (RRHO2) output plot. Genes from each list are ordered from most upregulated to most downregulated, with the most upregulated gene of each differential gene list in the lower left corner. b) RRHO2 plots comparing differentially expressed genes (DEGs) between control and *PTEN* heterozygous mutant organoids at one month *in vitro* versus genes changing over time in organoids between 23 days and 1.5 months *in vitro*, in the newborn DL PN and immature DL PN populations. c) Spearman correlation of DEGs between each cortical cell type within control and *PTEN* mutant organoids at one month, and genes that change within that cell type from 23 days to 1.5 months in control organoids (Uzquiano et al., bioRxiv, 2022). Correlation is calculated on each gene’s signed logFC. d) RRHO2 plots comparing differentially expressed genes between control and *PTEN* heterozygous mutant organoids at three month *in vitro* versus genes changing over time in organoids between two and four months *in vitro*, in the CFuPN and CPN populations. e) Spearman correlation of DEGs between each cortical cell type within control and *PTEN* heterozygous mutant organoids at three months, and genes that change within that cell type from two months to four months in control organoids (Uzquiano et al., bioRxiv, 2022). Correlation is calculated on each gene’s signed logFC. f) Volcano plot showing fold change versus FDR of measured proteins in MS experiments comparing *PTEN* heterozygous versus control organoids cultured for 35 days (n = 4 single organoids per genotype). Significant DEPs are shown in red (FDR<0.1). Proteins mentioned in the text are highlighted. g) Enriched gene ontology terms for DEPs between *PTEN* heterozygous and control organoids cultured for one month *in vitro*. h) Protein expression changes at two vs. one month for control and mutant organoids. Gray lines connect values for the same protein in the two genotypes. P-value from a paired signed Wilcoxon rank test. Only significant DEPs are shown; for the same analysis done with all detected proteins, see Supplementary Figure 3. i) Comparison of protein expression changes in *PTEN* heterozygous vs. control organoids at 1 month (bottom) versus changes in two vs. one month control organoids (top). Color and y-axis indicate log2 fold change. Only proteins with significantly differential expression between control and mutant at one month are shown. (n = 3 single organoids per genotype for 2 months). j) Schematic summarizing line-specific developmental acceleration/deceleration phenotypes from scRNA-seq and proteomics experiments. DL, deep layer; CPN, callosal projection neurons; CFuPN, corticofugal; CTL, control; FC, Fold change.

Strikingly, heterozygous *PTEN* mutation induced differences in predicted developmental speed across many organoid cell types, more prominently at three months *in vitro* (**Supplementary Fig. 3a,c**); that is, a significantly larger or smaller overlap between the genes up(down)-regulated in each cell population in the *PTEN* heterozygous mutant with the genes up(down)-regulated in the same cell population over time than expected by chance. Notably, the direction of this effect differed between donors: while most of the Mito210 cell types displayed gene expression differences indicating accelerated development, the PGP1 cell types were predominantly predicted to be delayed, suggesting at genetic context as an important modulator of the *PTEN* phenotype (**Supplementary Fig. 3**). Importantly, not all cells showed this effect, e.g., DEGs in heterozygous cortical hem and CPN did not show a correlation with developmentally-regulated genes, suggesting cell type specificity of this phenotype (**Supplementary Fig. 3a,c**). To confirm this overall trend, we performed bulk RNA-seq profiling on single *PTEN* heterozygous and isogenic control PGP1 organoids from an independent differentiation batch at one month. Consistent with the scRNA-seq results, DEGs upregulated in heterozygous relative to control organoids were correlated with developmentally-downregulated genes in the organoid atlas between 23 days and 1.5 months, confirming a predicted developmental delay in *PTEN* heterozygous organoids (**Supplementary Fig. 3b**).

Among projection neurons, in the one-month comparison, deep-layer projection neurons (DL PN) in the Mito210 heterozygous organoids showed gene expression changes suggesting accelerated differentiation in the mutant; by contrast, PGP1 heterozygous organoids indicated delayed differentiation of deep layer projection neurons at this age (**Fig. 2b,c**). Effects on deep-layer neuron populations were still evident at three months: again, Mito210 corticofugal projection neurons (CFuPN) appeared to be accelerated in their development, while PGP1 CFuPN were delayed (**Fig. 2d, e**). This effect was seen in all differentiation batches, at both ages (**Supplementary Fig. 3a,c**). Other neuronal populations such as CPN did not show significant association with developmental progression, indicating cell-type specificity among projection neurons. Thus, heterozygous *PTEN* mutation results in changes in predicted developmental timing in both the Mito210 and PGP1 donor cell lines, which is modulated by both cell type and by genetic background.

To examine the molecular changes induced by PTEN mutation at the protein level, we performed proteomic analysis of control and mutant organoids for the Mito210 line (**Fig. 2f-j** and **Supplementary Fig. 3d, e**). Whole-proteome mass spectrometry of organoids at one month in culture (n = 4 single organoids per genotype) showed significant differential expression of 75 proteins between genotypes (FDR<0.1, moderated t-test; **Fig. 2f**). Gene set enrichment analysis (GSEA) identified “synapse” as the most enriched biological process, along with exocytosis (e.g., STX1B) and the calcium signaling pathway (e.g., PPP3CB, CALM3, and CAMK4), pointing to a potential effect of the *PTEN* mutation on neuronal maturation (**Fig. 2g**). We also performed this whole-proteome analysis at a later time point, two months (n = 3 single organoids per genotype). While there were no differences in bulk protein abundance between mutant and control at 2 months, when we examined the proteins that were differentially expressed between time points (two months vs. one month) in each genotype, the mutant organoids showed significantly smaller expression changes over time than controls did (p = 1.6*10^-13^, Wilcoxin rank test, **Fig. 2h** and **Supplementary Fig. 3d**), indicating that the mutant 1 month organoids resembled their older counterparts more closely than the stage-matched control organoids did. Furthermore, the mutation-driven changes (i.e., the set of proteins that were differentially expressed between genotypes in the 1 month organoids) were correlated with time-driven changes (i.e., the set of proteins that changed in both the control and mutant proteomes through time, 2 months vs. 1 month), indicating that heterozygous loss of function of *PTEN* affects the expression of developmentally-regulated proteins (Pearson’s r = 0.76, p < 10^-15^, **Fig. 2i, j** and **Supplementary Fig. 3e**). This result points to a precocious development of Mito210 *PTEN* mutant organoids, consistent with the predicted developmental acceleration observed in the scRNA-seq data from this line.

In humans, different types of mutations and dosages of *PTEN* result in different diseases: heterozygous loss-of-function of *PTEN* has been associated with ASD (Busch et al., Transl Psychiatry, 2019; Butler et al., J Med Genet, 2005; Frazier et al., Mol Psychiatry, 2015, Serebriiskii et al., 2022), while both heterozygous and homozygous *PTEN* loss-of-function has been associated with tumor syndromes (Krohn et al., Am J Pathol, 2012; Wang et al., Clin Cancer Res, 1998). We therefore investigated whether different dosages of *PTEN* may result in different phenotypes also in brain organoids. To test this, we generated homozygous *PTEN* mutations in the PGP1 parental cell line, using the same gRNA to target both alleles of the gene (**Methods**, **Supplementary Fig. 1a**). We then generated cortical organoids with the same protocol as above (**Supplementary Fig. 4a**). Western Blot analysis performed in the homozygous line showed a complete absence of PTEN, accompanied by an increased phosphorylation of the downstream effectors AKT and ERK (MAPK1), which was much more pronounced than in the heterozygous line (**Supplementary Fig. 1c**).

Profiling of 19,944 single cells by scRNA-seq at one month (**Figure 3a**, Supplementary Figure 4b) showed that among the cortical cell types, the only significant change in cell type composition was a moderate decrease of IPC in *PTEN* homozygous mutant compared to control organoids (n = 3 single organoids per genotype, FDR=0.001, logistic mixed models; **Figure 3b** and **Supplementary Figure 4c, d**). Analysis of 11,337 single cells profiled from three month old homozygous mutant PGP1 organoids (**Fig. 3c** and **Supplementary Fig. 4e**) revealed an increase in the proportion of oRG (FDR=0.001), as previously observed in another organoid model grown from *PTEN* homozygous mutant iPSC (Li et al., Cell Stem Cell, 2017), and of aRG (FDR=0.001); these two populations were also increased in the heterozygous mutant (**Fig. 3d** and **Supplementary Fig. 4f, g**). This indicates that overproduction of the oRG population is a consistent outcome of reduced *PTEN*, in both homozygous and heterozygous settings, independent of genetic background. However, we observed differences between the heterozygous and homozygous mutants; assignment of cells to phases of the cell cycle based on expression of related genes revealed a decrease in the proportion of proliferating oRG (FDR=3.12*10^-15^, **Supplementary Fig. 4h**). In addition, the proportions of neuronal populations showed different effects in the *PTEN* homozygous mutant (**Fig. 3g-h**). Collectively, the data indicate that gene dosage affects the phenotypic manifestation of *PTEN* mutations (Trotman et al., 2003), underscoring the importance of studying this gene in the heterozygous state, similar to that found in ASD patients.

**Figure 3.**
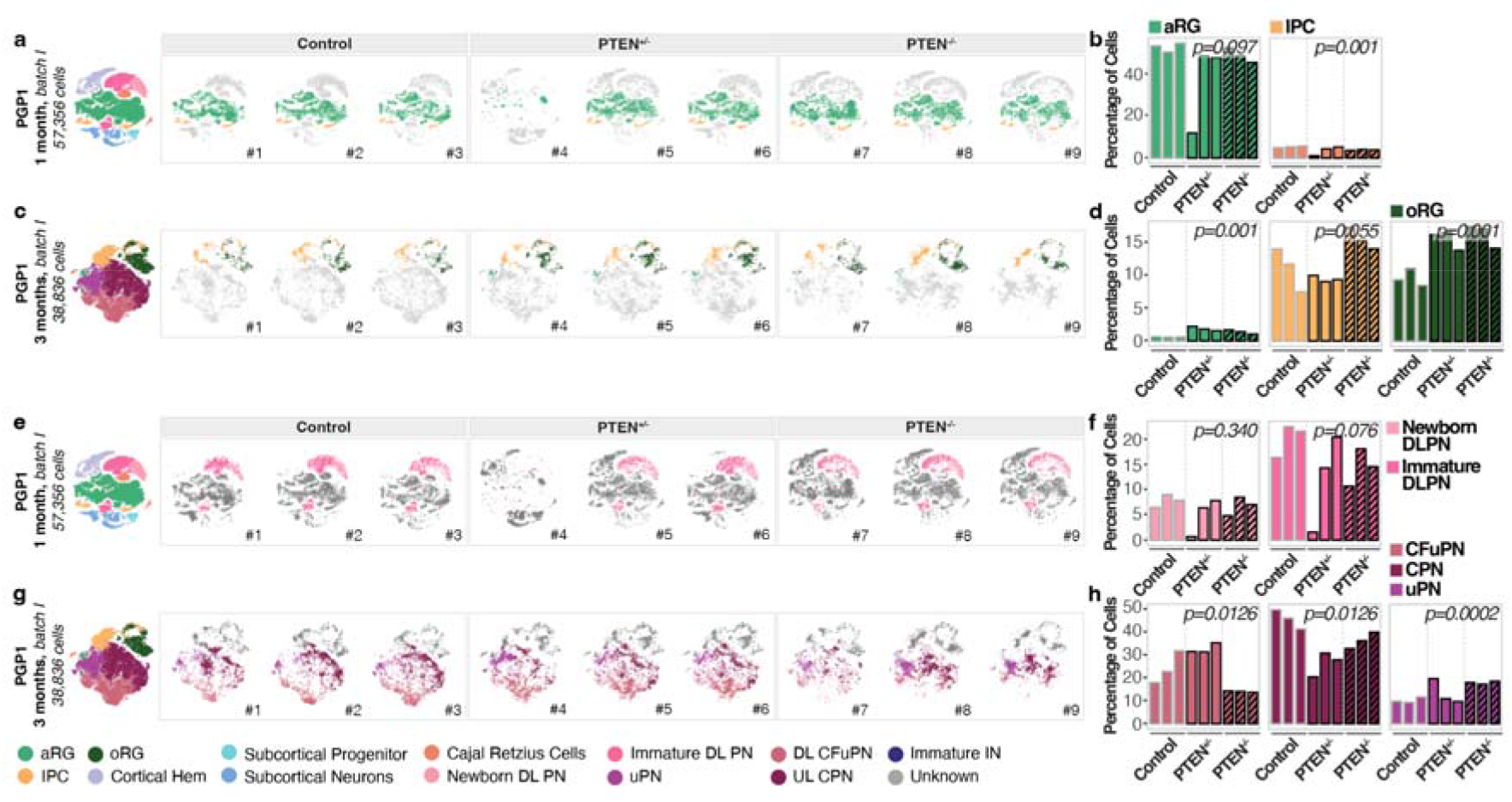
Differences in *PTEN* dosage result in convergent progenitor phenotypes, but divergent projection neurons abnormalities. a) Left: t-SNE plot of scRNA-seq data, color-coded by cell type, from one month PGP1 control, *PTEN* heterozygous and *PTEN* homozygous mutant organoids (n = 3 single organoids per genotype). Right: t-SNE plots for control, *PTEN* heterozygous, and *PTEN* homozygous mutant individual organoids, with cell types of interest highlighted in color: apical radial glia (aRG, light green), intermediate progenitors cells (IPC, yellow) and outer radial glia (oRG, dark green). b) Bar charts showing the percentage of cells for the highlighted cell populations in a, right, in each control, *PTEN* heterozygous, and *PTEN* homozygous mutant organoid at one month *in vitro*. FDRs for a difference in cell type proportions between control and mutant, based on logistic mixed models (see Methods) are shown. c) Left: t-SNE plot of scRNA-seq data, color-coded by cell type, from three months PGP1 control, *PTEN* heterozygous and *PTEN* homozygous mutant organoids (n = 3 single organoids per genotype). Right: t-SNE plots for control, *PTEN* heterozygous, and *PTEN* homozygous mutant individual organoids, with cell types of interest highlighted in color: apical radial glia (aRG, light green), intermediate progenitor cells (IPC, yellow) and outer radial glia (oRG, dark green). d) Bar charts showing the percentage of cells for the highlighted cell populations in c, right, in each control, *PTEN* heterozygous, and *PTEN* homozygous mutant organoid at three months *in vitro*. FDRs for a difference in cell type proportions between control and mutant, based on logistic mixed models (see Methods) are shown. e) Left: t-SNE plot of scRNA-seq data, color-coded by cell type, from one month PGP1 control, *PTEN* heterozygous and *PTEN* homozygous mutant organoids (n = 3 single organoids per genotype). Right: t-SNE plots for control, *PTEN* heterozygous, and *PTEN* homozygous mutant individual organoids, with cell types of interest highlighted in color: newborn deep layer projection neurons (Newborn DLPN, light pink), immature deep layer projection neurons (Immature DLPN, dark pink). f) Bar charts showing the percentage of cells for the highlighted cell populations in e, right, in each control, *PTEN* heterozygous, and *PTEN* homozygous mutant organoid at one month *in vitro*. FDRs for a difference in cell type proportions between control and mutant, based on logistic mixed models (see Methods) are shown. g) Left: t-SNE plot of scRNA-seq data, color-coded by cell type, from three months PGP1 control, *PTEN* heterozygous and *PTEN* homozygous mutant organoids (n = 3 single organoids per genotype). Right: t-SNE plots for control, *PTEN* heterozygous, and *PTEN* homozygous mutant individual organoids, with cell types of interest highlighted in color: corticofugal projection neurons (CFuPN, pink), callosal projection neurons (CPN, burgundy), unspecified projection neurons (uPN, purple). h) Bar charts showing the percentage of cells for the highlighted cell populations in g, right, in each control, *PTEN* heterozygous, and *PTEN* homozygous mutant organoid at one month *in vitro*. FDRs for a difference in cell type proportions between control and mutant, based on logistic mixed models (see Methods) are shown. aRG, apical radial glia; DL, deep layer; UL, upper layer; PN, projection neurons; oRG, outer radial glia; IPC, intermediate progenitor cells; CPN, callosal projection neurons; CFuPN, corticofugal projection neurons; uPN, unspecified PN; IN, interneurons; CTL, control.

**Figure 4.**
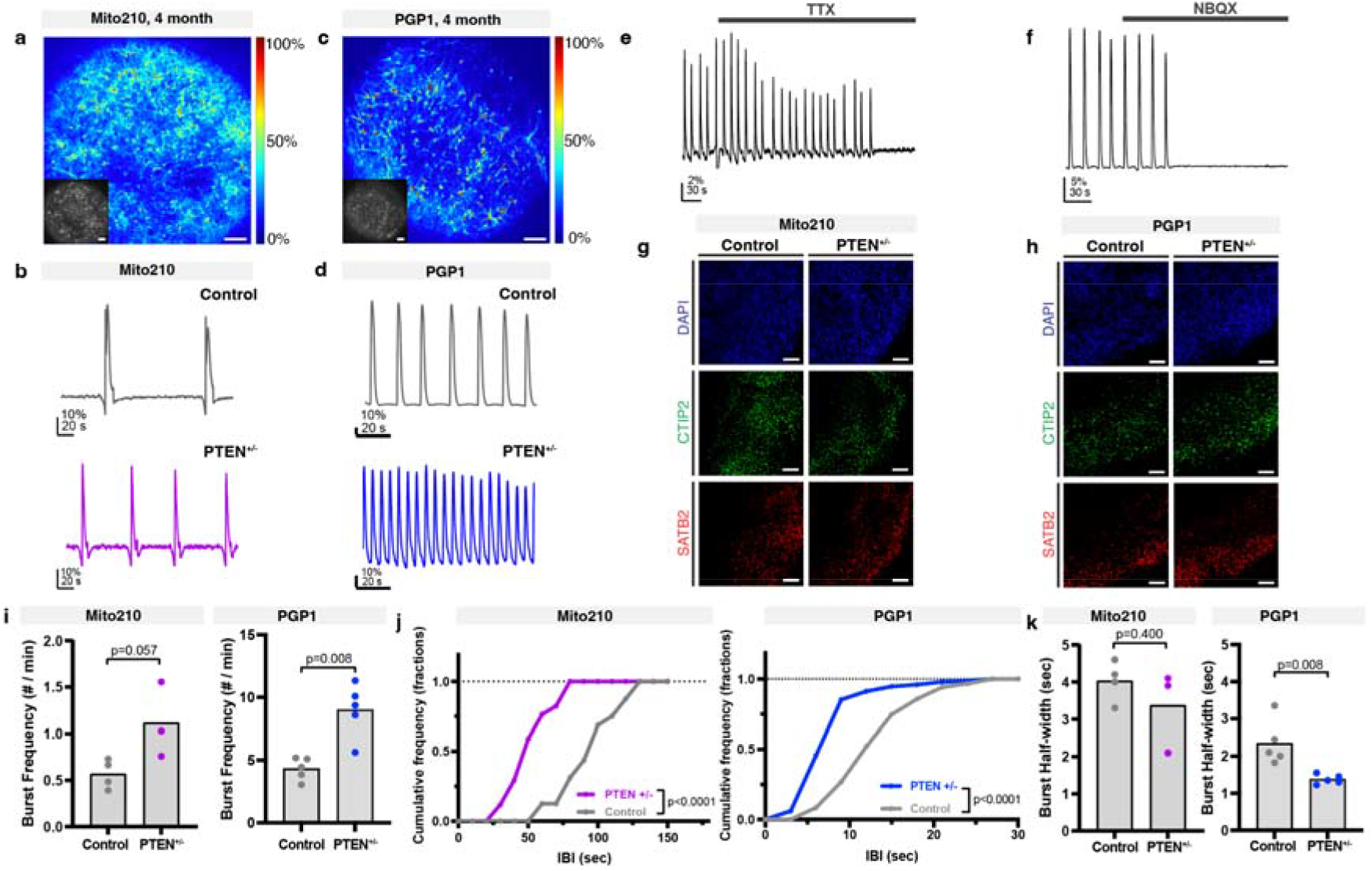
Developmental abnormalities in *PTEN* mutant organoids converge on altered circuit activity. a-d) Network bursting in cortical organoids. a, c) Representative images of intact Mito210 (a) and PGP1 (c) control organoid at four months transduced with SomaGCaMP6f2, showing the ΔF/F signal at the peak of a network burst using a pseudocolored scale. Inset: Raw fluorescence image. Scale bar: 100 μm. b, d) Representative population-averaged calcium transient for control (top) and heterozygous mutant (bottom) organoids generated from Mito210 (b) and PGP1 (d) donor lines. e, f) Example population-averaged calcium transients in control PGP1 organoids upon bath application of 2 μM TTX (e) and 20 μM NBQX (f). g, h) Immunohistochemistry for CTIP2 (green) and SATB2 (red) in control and *PTEN* heterozygous mutant organoids at three months, from both Mito210 (g) and PGP1 (h) cell lines. Scale bars: 100 μm. i) Spontaneous network burst frequency. The dots show the average values per organoid and the bars show the mean across all organoids. j) Cumulative frequency distribution of inter-burst interval (IBI) for control and mutant organoids. k) Spontaneous network burst duration. The dots show the average values per organoid and the bars show the mean across all organoids. Min, minutes; sec, seconds.

Correct formation and function of the cortical local circuit depends on the timely integration and maturation of multiple neuronal subtypes. We hypothesized that the asynchronous development of specific neuronal populations seen in the *PTEN* heterozygous mutant organoids may affect the function of cortical local circuits. To test this, we performed calcium imaging analysis in intact preparations of four-month-old heterozygous mutant and control organoids from both genetic backgrounds. We analyzed spontaneous neuronal activity by transducing organoids with adeno-associated viruses bearing the calcium indicator CAG::SomaGCaMP6f2 (Shemesh et al., Neuron, 2020) (**Methods**), and performing acute recordings of intracellular calcium dynamics (**Fig. 4**). Both control and mutant organoids displayed spontaneous activity (163 +/− 77 active cells from 183 +/− 79 total cells, per imaged section, **Fig. 4a-d**), with the main form of activity being network bursting (**Fig. 4a-d**). When organoids were treated with the voltage-gated sodium channel blocker tetrodotoxin (TTX), calcium transients were abolished, suggesting that activity was mediated by action potentials (**Fig. 4e**, **Supplementary Fig. 5a**). To verify the participation of excitatory neurons (**Fig. 4f-h**), we treated organoids with an antagonist of non-NMDA glutamate receptors, 2,3-dioxo-6-nitro-1,2,3,4-tetrahydorbenzo[*f*]quinoxaline-7-sulfonamide (NBQX), which suppressed spontaneous activity, indicating that synchronized network bursting is driven by functional glutamatergic synapses (**Fig. 4f**, **Supplementary Fig. 5b**).

*PTEN* heterozygous organoids showed an increased level of bursting activity compared to control, in both the Mito210 and PGP1 genetic backgrounds (Mito210 burst frequency p=0.057, IBI cumulative frequency p<0.0001, 4 controls vs 3 heterozygous, Mann-Whitney test and Kolmogorov-Smirnov test, respectively; PGP1 burst frequency p=0.008, IBI cumulative frequency p<0.0001, n = 5 organoids per condition, Mann-Whitney test and Kolmogorov-Smirnov test, respectively, **Fig. 4i, j**). In addition, bursts had shorter duration in the heterozygous organoids compared to controls in the PGP1 background (p = 0.008, n = 5 organoids per condition, Mann-Whitney test), and Mito210 organoids followed a similar trend (**Fig. 4k**). Interestingly, control organoids in the two parental lines displayed a striking difference in the level of spontaneous activity (burst per minute: 0.57 ± 0.16, Mito210 vs 4.33 ± 0.90, PGP1), suggesting that the degree of spontaneous neural activity may be an idiosyncratic trait that varies between donor lines. However, both backgrounds showed the same relative phenotype in the mutant (**Fig. 4i**), suggesting that these differences in genetic background do not dominate the mutant phenotype. We previously reported that bursting activity (burst frequency and burst half-width) is altered in organoids with a heterozygous mutation for a different ASD-associated gene, *SUV420H1* (Paulsen et al., Nature, 2022). Interestingly, although both genes affect bursting behavior, they do so in opposite directions, suggesting that mutations in different ASD risk genes may affect cortical circuit activity in different ways. Taken together, the data support the hypothesis that early abnormalities in the development and maturation of specific populations of neurons converge on later defects in circuit physiology, resulting in higher-order phenotypes that are shared across both genetic backgrounds and individual ASD risk genes.

Human brain organoids offer an unprecedented opportunity to investigate early phenotypic manifestations associated with risk genes for complex neurodevelopmental disorders, including ASD. Here, we investigated the function of *PTEN*, a gene previously associated with ASD (Sanders et al., Neuron, 2015; Satterstrom et al., Cell, 2020; Stessman et al., Nat Genet, 2017), in early stages of human cortical development. We reveal pleiotropic effects of *PTEN* mutation involving different steps of development, including alteration of specific progenitor populations (e.g., oRG), asynchronous development of deep layer projection neurons, and abnormal circuit activity.

While the oRG phenotype agrees with previously reported results in human brain organoids with homozygous loss-of-function of *PTEN* (Li et al., Cell Stem Cell, 2017), we found a previously undescribed effect of *PTEN* on human projection neuron development. Interestingly, while the progenitor phenotype is independent of *PTEN* dosage, changes in neuronal proportion were seen primarily in the homozygous mutant. These results point to the importance of studying risk genes in contexts that recapitulate the gene dosage state found in patients.

Different human genomic contexts have been suggested to modulate the phenotypic expressivity of ASD-risk genes in a mutation- and phenotype-dependent manner (Paulsen et al., Nature, 2022). Here we further confirm the importance of investigating ASD-risk mutations across different genomic contexts; we find that the same *PTEN* mutation affects the development of the same neuronal population in lines from different donors, but it does so in opposite directions in the two genetic backgrounds. These results point towards mechanisms that may underlie the variability of clinical manifestations observed in human patients with mutation in the same risk gene. It is intriguing that, despite the divergent molecular and cellular phenotypes observed for different ASD risk genes and in different genetic backgrounds, these effects ultimately may converge on changes in local circuit activity. This supports the hypothesis that the effects of different ASD-risk genes may converge on similar features of higher-order processes of circuit activity and function.

## Data availability

See our interactive portal to explore the single cell RNA-seq data at https://singlecell.broadinstitute.org/single_cell/study/SCP1964. Full datasets will be available for download after publication.

## Author contributions

P.A., M.P., A.U. and B.P, conceived the experiments. M.P., R.S. and A.T, generated, cultured and characterized all organoids and P.A. supervised their work. X.A. performed bulk and scRNA-seq experiments with help from M.P., A.U. R.S. and S.V.; X.J. and E.M. performed Slide-seq experiments, with the supervision of F.C.; A.J.K. and J.Z.L. performed scRNA-seq analysis and J.Z.L. and A.R. supervised the work. K.K. performed the bulk RNA-seq analysis. M.P., A.U., B.P., S.V. and A.J.K. worked on cell type assignments and data analysis. K.T., M.P., A.J.K. performed proteomics analysis, supervised by K.L.; S.M.Y and P.S. performed the calcium imaging experiments and analysis, supervised by E.S.B and with the help of R.S.; L.B. generated the Mito210 *PTEN* mutant line; P.A., M.P., A.U., B.P. and A.J.K. wrote the manuscript with contributions from all authors. All authors read and approved the final manuscript.

## Acknowledgments

We thank J. R. Brown for editing the manuscript, and the entire Arlotta Lab for support and insightful discussions; V. Vuong, C. Abbate and S.N. Smith for technical support in organoid culture and characterization; N. Haywood for scRNA-seq experiments; A. Shetty for help with scRNA-seq cell type classifications; S. Simmons for revising the manuscript; the Broad Genomics Platform for sequencing; the Cohen laboratory for the Mito 210 line; L.M. Daheron at the Harvard Stem Cell Facility for expanding edited lines and generating the PGP1 *PTEN* mutant lines; A. Podury for helping to establish the calcium imaging protocols; and B. Budnik at the Harvard Center for Mass Spectrometry for conducting proteomics experiments. This work was supported by grants from the Stanley Center for Psychiatric Research, the Broad Institute of MIT and Harvard, the National Institutes of Health (R01MH112940 to P.A. and J.Z.L., and P50MH094271, U01MH115727, and RF1MH123977 to P.A.), the Klarman Cell Observatory to J.Z.L. and A.R., and the Howard Hughes Medical Institute to A.R. AR was a Howard Hughes Medical Institute and a Koch Institute extramural member while conducting this study.

## Conflicts of Interests

P.A. is a SAB member at Herophilus, Rumi Therapeutics, and Foresite Labs, and is a co-founder of Vesalius. A.R. is a founder and equity holder of Celsius Therapeutics, an equity holder in Immunitas Therapeutics and until August 31, 2020 was a SAB member of Syros Pharmaceuticals, Neogene Therapeutics, Asimov and Thermo Fisher Scientific. From August 1, 2020, A.R. is an employee of Genentech and has equity in Roche. From September 1, 2021, M.P. is an employee of Roche.

## Supplementary Figures

**Supplementary Figure 1 (Figure 1).**
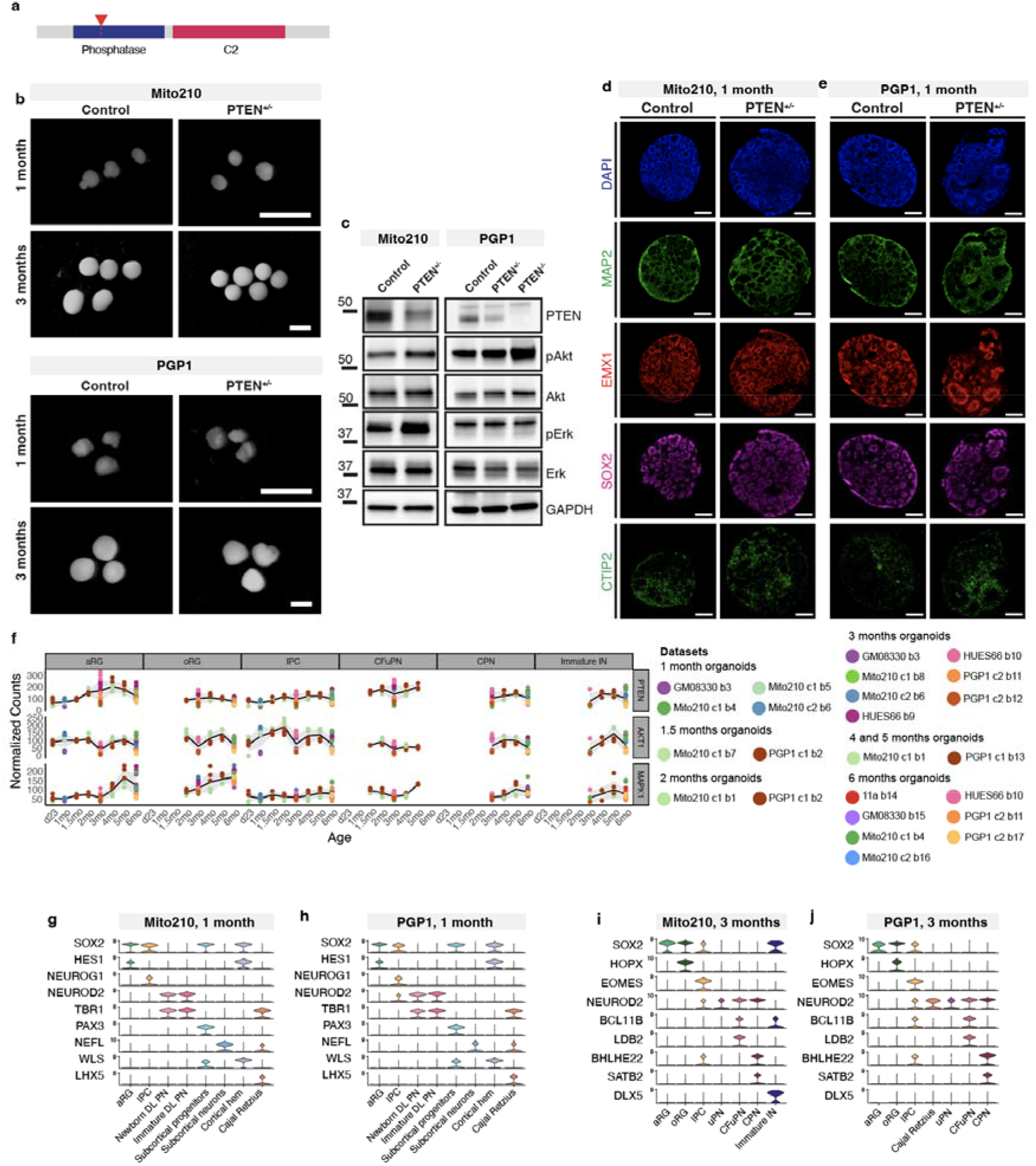
a) Protein domain structure of PTEN. The Mito210 and PGP1 parental lines were edited using CRISPR-Cas9 to introduce a protein-truncating mutation in the phosphatase domain (red triangle). b) Photomicrograph of intact Mito210 (top) and PGP1 (bottom) *PTEN* heterozygous mutant and control organoids at one and three months. Scale bars: 3 mm. c) Western blot analysis showing PTEN protein expression in control lines, and its reduction in the mutant lines, as well as both total and phosphorylated AKT and ERK (downstream effectors of PTEN) in control and *PTEN*^+/−^ lines. Molecular weight in kDa is shown on the left of the gel. d, e) Immunohistochemistry for neuronal (MAP2), dorsal forebrain neural progenitor (EMX1, SOX2), and CFuPN (CTIP2) markers in one month organoids derived from the Mito210 *PTEN* heterozygous mutant and isogenic control cell lines (e) and PGP1 *PTEN* heterozygous mutant and isogenic control cell lines (f). Scale bar, 200 μm. f) Gene expression (as normalized read counts) of *PTEN, AKT* and *MAPK1* across ambient-RNA-corrected gene expression throughout organoid development, 23 days to 6 months *in vitro* (Uzquiano et al. 2022). Each colored dot represents the normalized counts within a differentiation batch. g-h) Violin plots showing marker gene expression across cell types in control and *PTEN* heterozygous mutant organoids cell from the Mito210 (g) and PGP1 (h) cell lines at one month *in vitro*. Data were generated using datasets shown in Figure 1a and b, left. i-j) Violin plots showing marker gene expression across cell types in control and *PTEN* heterozygous mutant organoids cell from the Mito210 (i) and PGP1 (j) cell lines at three months *in vitro*. Data were generated using datasets shown in Figure 1c and d, left. aRG, apical radial glia; DL, deep layer; UL, upper layer; PN, projection neurons; oRG, outer radial glia; IPC, intermediate progenitor cells; CPN, callosal projection neurons; CFuPN, corticofugal projection neurons; uPN, unspecified PN; IN, interneurons.

**Supplementary Figure 2 (Figure 1).**
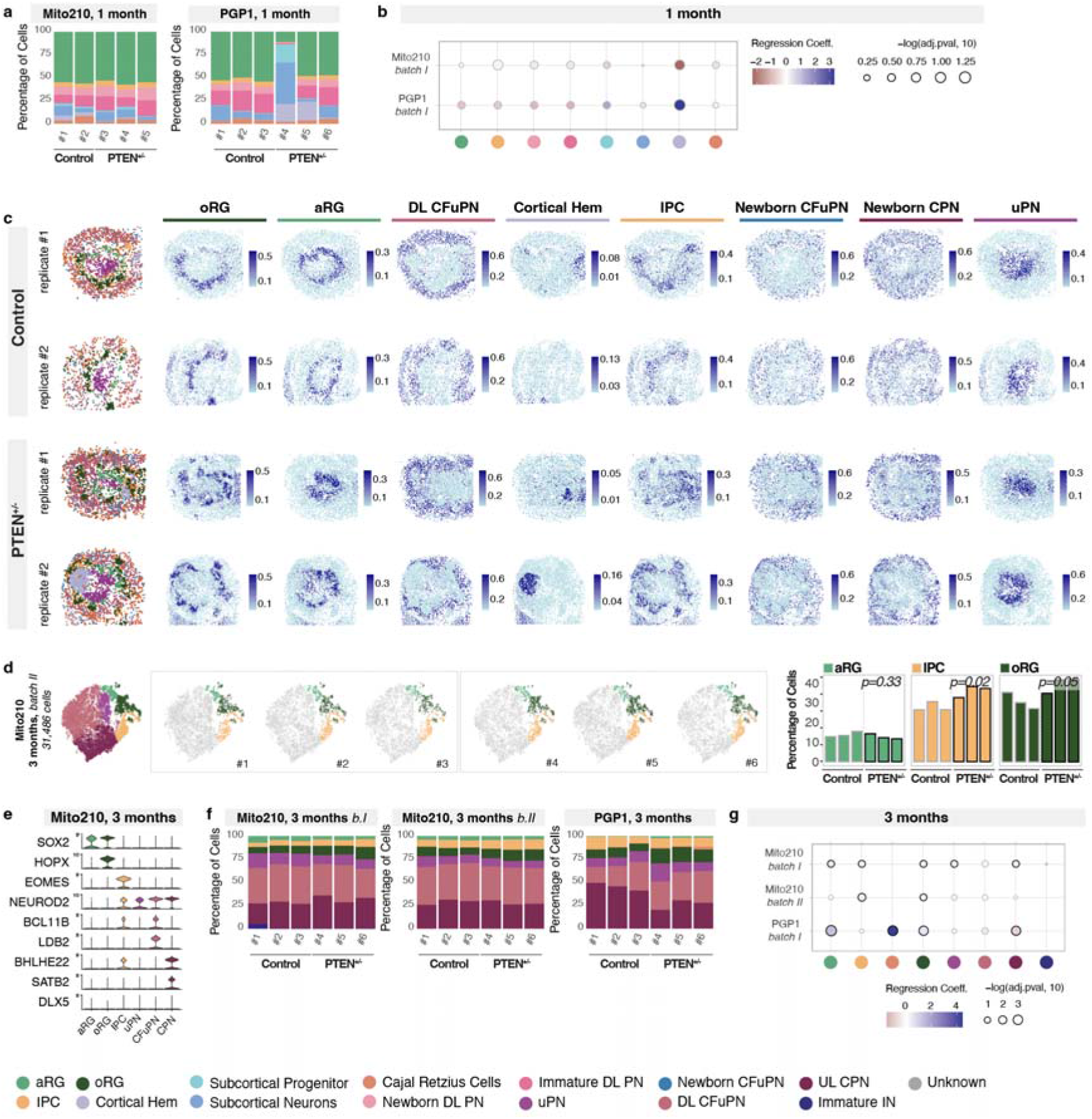
a) Barcharts showing the percentage of cells for all the cell populations in each control and *PTEN* heterozygous mutant organoid at one month. b) Changes in cell type composition between control and *PTEN* heterozygous mutant organoids at one month. For each cell type and differentiation batch, points are colored by the effect size of a change in cell type proportion (blue indicates an increase in that cell type in mutant, red indicates a decrease), and sized by the statistical significance of the change. Both calculated using logistic mixed models (see Methods). The largest effect on non-cortical cell types was seen in the cortical hem population; however, the direction of effect was inconsistent between cell lines, and previous data showed a higher-than-average variability in the proportion of this population across differentiation batches in control organoids (Uzquiano et. al., 2022), so conclusions regarding the number of cells in the cortical hem population of these organoids should be made with caution. c) Spatial plots of Slide-seqV2 data from 2 control and 2 *PTEN* mutant organoids at two months. Left: Spatial plots colored by RCTD-assigned cell type. Right, spatial plots showing RCTD prediction weights. d) Left: t-SNE plot of scRNA-seq data, color-coded by cell type, from Mito210 batch II three months control and *PTEN* heterozygous mutant organoids (Mito210 control n = 3, Mito210 heterozygous n = 3; see Figure 1a for batch I). Middle: t-SNE plots for control and mutant individual organoids, with cell types of interest highlighted in color: apical radial glia (aRG, light green), intermediate progenitor cells (IPC, yellow) and outer radial glia (oRG, dark green). Right: bar charts showing the percentage of cells for the highlighted cell populations in each control and mutant organoid. FDRs for a difference in cell type proportions between control and mutant, based on logistic mixed models (see Methods) are shown. e) Violin plots showing marker gene expression (normalized counts) across cell types in control and *PTEN* heterozygous mutant organoids cell from the Mito210 cell line batch II at three months *in vitro*. Data were generated using datasets shown in (d). f) Barcharts showing the percentage of cells for all the cell populations in each control and *PTEN* heterozygous mutant organoid at three months. g) Changes in cell type composition between control and *PTEN* heterozygous mutant organoids at three months. aRG, apical radial glia; DL, deep layer; UL, upper layer; PN, projection neurons; oRG, outer radial glia; IPC, intermediate progenitor cells; CPN, callosal projection neurons; CFuPN, corticofugal projection neurons; uPN, unspecified PN; IN, interneurons.

**Supplementary Figure 3 (Figure 2).**
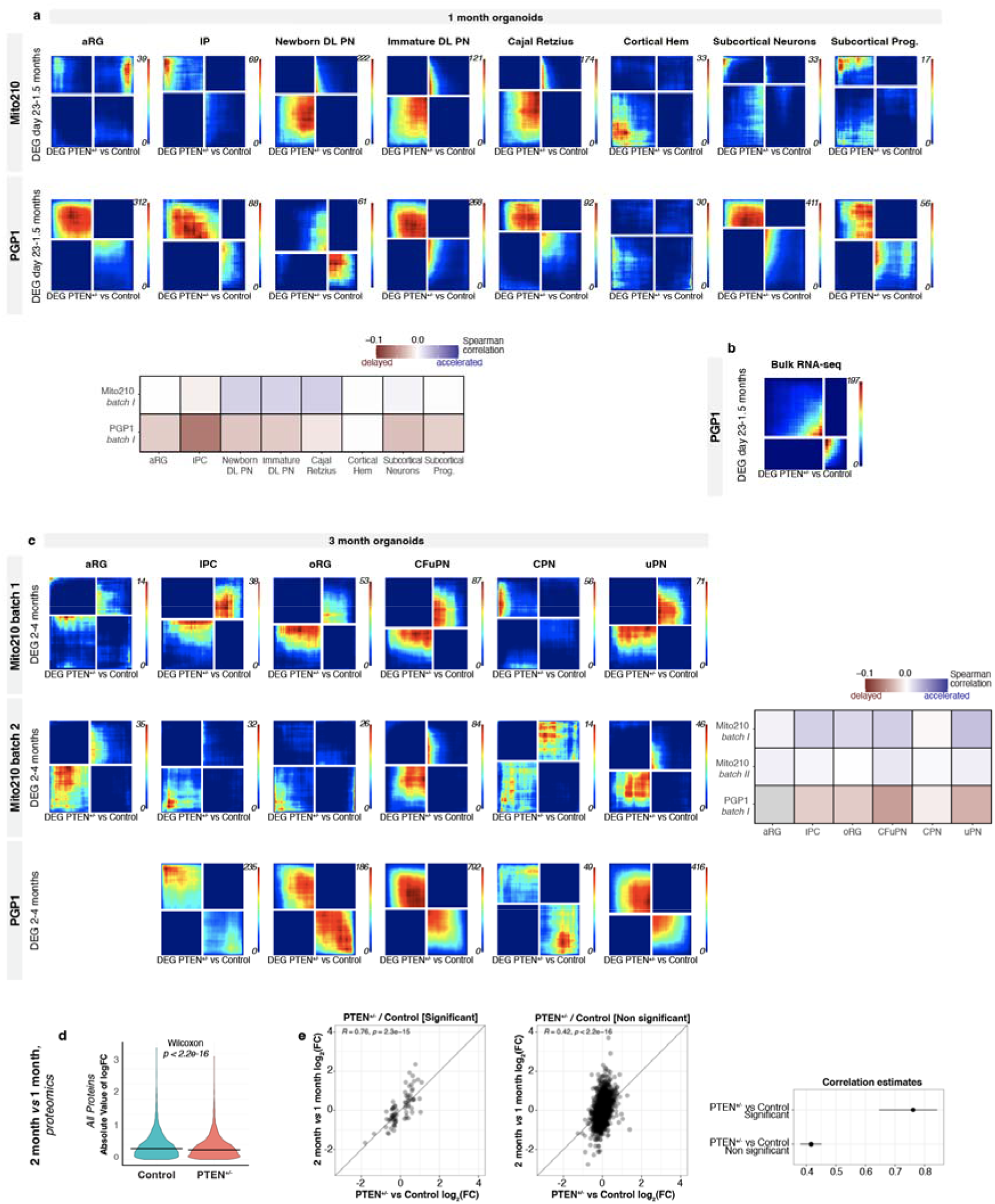
a) Top: RRHO2 plots comparing differentially expressed genes (DEGs) between control and *PTEN* heterozygous mutant organoids at one month *in vitro* versus genes changing over time in organoids between 23 days and 1.5 months *in vitro* across all cell types. Bottom: Spearman correlation of DEGs between each cell type within control and *PTEN* mutant organoids at one month, and genes that change within that cell type from 23 days to 1.5 months in control organoids (Uzquiano et al., bioRxiv, 2022). Correlation is calculated on each gene’s signed logFC. b) RRHO2 plots comparing differentially expressed genes (DEGs) between control and *PTEN* heterozygous mutant organoids at one month *in vitro* from bulk RNA-seq versus genes changing over time in organoids between 23 days and 1.5 months *in vitro* across all cell types. c) Top: RRHO2 plots from comparing differentially expressed genes between control and *PTEN* homozygous mutant organoids at three month *in vitro* and genes changing over time in organoids between two and four months *in vitro* across all cell types. Bottom: Spearman correlation of DEGs between each cortical cell type within control and *PTEN* mutant homozygous organoids at three months, and genes that change within that cell type from 2 months to 4 months in control organoids (Uzquiano et al., bioRxiv, 2022). Correlation is calculated on each gene’s signed logFC. d) Protein expression changes between 2 and 1 month for control and mutant organoids, for all detected proteins. P-value from a paired signed Wilcoxon rank test. e) Correlation between developmentally-driven protein expression differences (control organoids at 2 versus 1 month) and mutation-induced protein expression differences (*PTEN* heterozygous versus control organoids at 1 month). Correlation is shown separately for proteins that showed a significant change between mutant and control organoids at 1 month (middle), and those that did not (right). Left, comparison of the respective Pearson correlation coefficients. aRG, apical radial glia; DL, deep layer; UL, upper layer; PN, projection neurons; oRG, outer radial glia; IPC, intermediate progenitor cells; CPN, callosal projection neurons; CFuPN, corticofugal projection neurons; uPN, unspecified PN; IN, interneurons; prog, progenitors.

**Supplementary Figure 4 (Figure 3).**
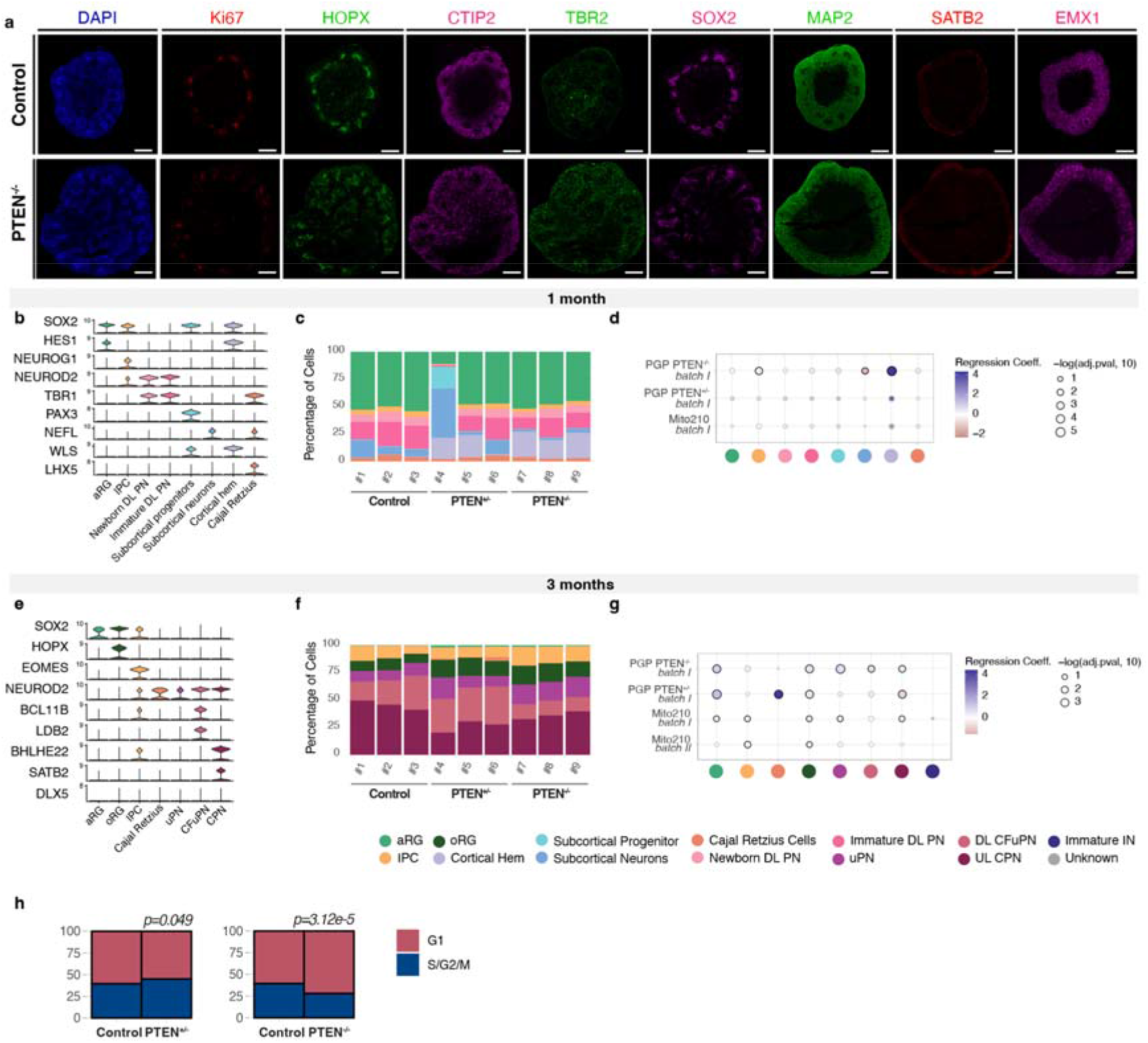
a) Immunohistochemistry for markers of dorsal forebrain neural progenitors (EMX1, SOX2), proliferating progenitors (Ki67), intermediate progenitor cells (TBR2), outer radial glia (HOPX), neurons (MAP2), CFuPN (CTIP2), and CPN (SATB2), in PGP1 *PTEN* homozygous mutant and isogenic control organoids at two months. Scale bar, 500 μm. b) Violin plots showing marker gene expression across cell types in control, *PTEN* heterozygous, and *PTEN* homozygous mutant organoids cell from the PGP1 cell line at one month *in vitro*. Data were generated using datasets shown in Figure 3a, e, left. c) Barcharts showing the percentage of cells for all the cell populations in each control, *PTEN* heterozygous, and *PTEN* homozygous mutant organoid at one month. d) Changes in cell type composition between control, *PTEN* heterozygous, and *PTEN* homozygous mutant organoids at one month. For each cell type and differentiation batch, points are colored by the effect size of a change in cell type proportion (blue indicates an increase in that cell type in mutant, red indicates a decrease), and sized by the statistical significance of the change. Both calculated using logistic mixed models (see Methods). e) Violin plots showing marker gene expression across cell types in control, *PTEN* heterozygous, and *PTEN* homozygous mutant organoids cell from the PGP1 cell line at three months *in vitro*. Data were generated using datasets shown in Figure 3c, g, left. f) Barcharts showing the percentage of cells for all the cell populations in each control, *PTEN* heterozygous, and *PTEN* homozygous mutant organoid at three months. g) Changes in cell type composition between control, *PTEN* heterozygous, and *PTEN* homozygous mutant organoids at three months. For each cell type and differentiation batch, points are colored by the effect size of a change in cell type proportion (blue indicates an increase in that cell type in mutant, red indicates a decrease), and sized by the statistical significance of the change. Both calculated using logistic mixed models (see Methods). h) Cell cycle analysis of oRG, comparing the proportion of oRG predicted to be proliferating versus in G1 (see Methods) in control vs *PTEN^+/−^* mutant organoids (left) and control vs *PTEN* homozygous mutant organoids (right). aRG, apical radial glia; DL, deep layer; UL, upper layer; PN, projection neurons; oRG, outer radial glia; IPC, intermediate progenitor cells; CPN, callosal projection neurons; CFuPN, corticofugal projection neurons; uPN, unspecified PN; IN, interneurons.

**Supplementary Figure 5 (Figure 4).**
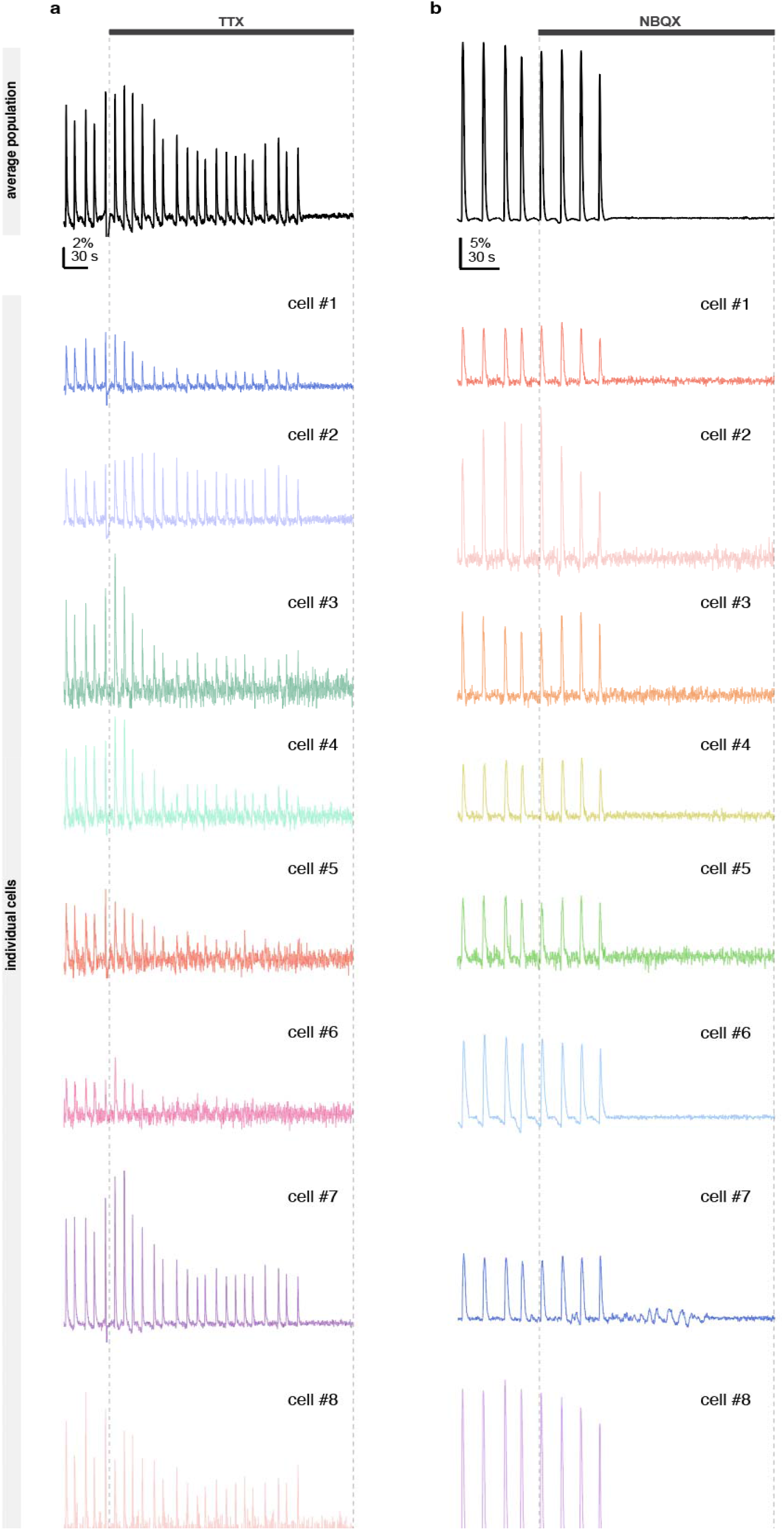
a, b) Representative calcium transients in control PGP1 organoids upon bath application of 2 μM TTX (a) and 20 μM NBQX (b); corresponding to Figure 4e and Figure 4f, respectively. The population-averaged signals are presented at the top, and individual traces for eight different cells per condition are displayed underneath the population transient. The experiments were performed for both TTX (four Mito210 and four PGP1 organoids) and NBQX (two Mito210 and five PGP1 organoids).

## Materials and methods

### Pluripotent stem cell culture

All experiments involving human cells were performed according to ISSCR 2016 guidelines, and approved by the Harvard University IRB and ESCRO committees. The Mito210 iPSC line was derived by B. Cohen Lab (McLean Hospital), and the PGP1 iPSC line by G. Church (Harvard University) (Church, Mol Syst Biol, 2005). Both Mito210 and PGP1 are human male iPSC lines from individuals with no known history of ASD or other psychiatric condition (Mito210 confirmed by structured psychiatric interview, PGP1 with publicly available records). Cell lines were cultured as previously described (Velasco et al., Nature, 2019; Velasco et al., Protocol Exchange, 2019). All human iPSC cultures were maintained below passage 50, were negative for mycoplasma (assayed with MycoAlert PLUS Mycoplasma Detection Kit, Lonza), and karyotypically normal (G-banded karyotype test performed by WiCell Research Institute). The Mito210 line was authenticated by genotyping analysis (Fluidigm FPV5 chip) performed by the Broad Institute Genomics Platform (in 2017). The PGP1 line was authenticated using STR analysis completed by TRIPath (in 2018).

### CRISPR guide design

The CRISPR guide for *PTEN* was designed using the Benchling CRISPR Guide Design Tool (Benchling Biology Software). The guide was designed to maximize on-target efficiency and minimize off-target sites in intragenic regions. It was chosen to induce a stop codon in the catalytic phosphatase domain of *PTEN* (gRNA: 5’-CCAAATTTAATTGCAGAGGT-3’ (AGG)).

### CRISPR-mediated gene editing

The Mito210 *PTEN* edited line was generated by the Broad Institute Stem Cell Facility. The guide RNA and Cas9 (EnGen Cas9 NLS from New England Biolabs) were transfected by using the NEON transfection system (Thermo Fisher Scientific, 1050 V, 30 ms, 2 pulses and 2.5×10^5^ cells).

The PGP1 *PTEN* mutant line was generated in collaboration with the Harvard Stem Cell Institute (HSCI) iPS Core Facility. Briefly, parental cells were transfected with the Neon system. For 100,000 cells 6.25 pmol TrueCut TM Cas9 Protein v2 (Thermo Fisher Cat: A36496) and 12.5 pmol of sgRNA (Synthego) were used. Post transfection, the pools of cells were harvested to test knock-out efficiency. The best pool was then selected for low density plating. A week later, colonies were picked and clones were screened by PCR and Sanger sequencing. Heterozygous and homozygous clones were expanded and the genotypes were re-confirmed post expansion.

### Sequence confirmation of edits

Insertions/deletions in individual clones were screened via PCR amplification using primers flanking the guide. The Mito210 *PTEN* mutant clone has a heterozygous 1 nucleotide insertion. The PGP1 *PTEN* mutant clone has a heterozygous 1 nucleotide insertion. The PGP1 *PTEN* homozygous clone has a 1 nucleotide insertion in one allele and 11 nucleotide insertion in the second allele.

### Organoid Differentiation

Dorsally patterned forebrain organoids were generated as previously described in (Velasco et al., 2019; Velasco, Paulsen and Arlotta, 2019). Briefly, on day 0, human iPSC or ESC, were dissociated to single cells with Accutase (Gibco), and 9,000 cells per well were reaggregated in ultra-low cell-adhesion 96-well plates with V-bottomed conical wells (sBio PrimeSurface plate; Sumitomo Bakelite) in Cortical Differentiation Medium (CDM) I, containing Glasgow-MEM (Gibco), 20% Knockout Serum Replacement (Gibco), 0.1 mM Minimum Essential Medium non-essential amino acids (MEM-NEAA) (Gibco), 1 mM pyruvate (Gibco), 0.1 mM 2-mercaptoethanol (Gibco), 100 U/mL penicillin (ThermoFisher), and 100 μg/mL streptomycin (Corning). For the PGP1 line cells were plated in the same pluripotent medium in which they were maintained for 1 day to better enable embryoid body formation. From day 0-6, ROCK inhibitor Y-27632 (Millipore) was added (final concentration of 20 μM). From day 0/1-18, Wnt inhibitor IWR1 (Calbiochem) and TGFβ inhibitor SB431542 (Stem Cell Technologies) were added (final concentration of 3 μM and 5 μM, respectively). From day 18, the aggregates were cultured in ultra-low attachment culture dishes (Corning) under orbital agitation in CDM II, containing DMEM/F12 medium (Gibco), 2mM Glutamax (Gibco), 1% N2 (Gibco), 1% Chemically Defined Lipid Concentrate (Gibco), 0.25 μg/mL fungizone (Gibco), 100 U/mL penicillin, and 100 μg/mL streptomycin. On day 35, cell aggregates were transferred to spinner-flask bioreactors (Corning) and maintained in CDM III (CDM II supplemented with 10% fetal bovine serum (FBS) (GE-Healthcare), 5 μg/mL heparin (Sigma) and 1% Matrigel (Corning)). From day 70, organoids were cultured in CDM IV (CDM III supplemented with B27 supplement (Gibco) and 2% Matrigel).

### Immunohistochemistry

Samples were fixed in 4% paraformaldehyde (PFA) (Electron Microscopy Services) for either 30 minutes (one month old organoids) or 1-3 hours (two month and older organoids). Samples were washed with 1X phosphate buffered saline (PBS) (Gibco), cryoprotected in a 30% sucrose solution overnight at 4□, embedded in optimum cutting temperature (OCT) compound (Tissue Tek), and cryosectioned at 14-18 μm thickness. Sections were washed with 0.1% Tween-20 (Sigma) in PBS, blocked for 1 hour at room temperature (RT) with 6% donkey serum (Sigma) + 0.3% Triton X-100 (Sigma) in PBS and incubated with primary antibodies overnight diluted with 2.5% donkey serum + 0.1% Triton X-100 in PBS. After washing, sections were incubated at room temperature with secondary antibodies diluted in the same solution as with primary antibodies (1:1000-1:1200) for 2 hours at room temperature or overnight at 4□, washed, and incubated with DAPI staining (1:10,000 in PBS + 0.1% Tween-20) for 15 minutes to visualize cell nuclei. Slices were mounted using Fluoromount-G (Invitrogen).

### Microscopy

Immunofluorescence images were acquired with the Lionheart™ FX Automated Microscope (BioTek Instruments), or with an Axio Imager.Z2 (Zeiss), and analyzed with the Gen5 (BioTek Instruments) or Zen Blue (Zeiss) image processing software.

### Western blotting

Proteins were extracted from iPSC using N-PER™ Neuronal Protein Extraction Reagent (Thermo Fisher Scientific) supplemented with protease (cOmplete™ Mini Protease Inhibitor Cocktail, Roche) and phosphatase inhibitor (PhosSTOP, Sigma). Lysates were centrifuged for 10 minutes at 13,500 rpm at 4° C. Supernatants were transferred to new tubes. Protein concentration was quantified using the Pierce™ BCA Protein Assay Kit (Thermo Fisher Scientific). 15-20 μg of protein lysates were separated on a NuPAGE™ 4-12%, Bis-Tris Gel (Invitrogen) or Mini-PROTEAN 4–15% Gels (Bio-Rad) and transferred onto PVDF membrane. Blots were blocked in 5% nonfat dry milk (Bio-Rad) and incubated with primary antibodies overnight. Afterward, the blots were washed and incubated at RT with secondary horseradish peroxidase-conjugated antibodies (Abcam) for 1 hour. Blots were developed using SuperSignal™ West Femto Maximum Sensitivity Substrate (Thermo Fisher Scientific) or ECL™ Prime Western Blotting System (Millipore), and ChemiDoc System (Bio-Rad).

### Dissociation of brain organoids and scRNA-seq

Individual brain organoids were dissociated into a single-cell suspension using the Worthington Papain Dissociation System kit (Worthington Biochemical). A detailed description of the dissociation protocol is available at Protocol Exchange, with adaptations depending on age and size (Quadrato, Sherwood and Arlotta, 2017; Velasco, Paulsen and Arlotta, 2019). We resuspended dissociated cells in ice-cold PBS containing 0.04% BSA (Sigma, PN-B8667), counted them with Cellometer K2 (Nexcelom Bioscience). Cells were loaded onto a Chromium™ Single Cell B or G Chip (10x Genomics, PN-1000153, PN-1000120), and processed through the Chromium Controller to generate single cell GEMs (Gel Beads in Emulsion). scRNA-seq libraries were generated with the Chromium™ Single Cell 3’ Library & Gel Bead Kit v3 or v3.1 (10x Genomics, PN-1000075, PN-1000121). We pooled libraries from different samples based on molar concentrations and sequenced them on a NextSeq 500 or NovaSeq instrument (Illumina) with 28 bases for read 1, 55 bases for read 2 and 8 bases for Index 1. If necessary, after the first round of sequencing, we repooled libraries based on the actual number of cells in each and re-sequenced with the goal of producing approximately 20,000 reads per cell for each sample.

### scRNA-seq data analysis

Reads from scRNA-seq were aligned to the GRCh38 human reference genome and cell-by-gene count matrices were produced with the Cell Ranger pipeline (v3.0.2, 10x Genomics) (Zheng et al., Nat Commun, 2017). Default parameters were used, except for the –cells’ argument. In Mito210 one month organoids, a control organoid was excluded because not enough cells could be recovered from that 10x channel. Data were analyzed using the Seurat R package v3.2.2 (Stuart et al., Cell, 2019) using R v3.6. Cell profiles expressing at least 500 genes were kept, and UM I counts were normalized for each cell by the total expression, multiplied by 10^6^, and log-transformed. Variable genes were found using the “mean.var.plot” method, and the ScaleData function was used to regress out variation due to differences in total UMIs per cell. Principal component analysis (PCA) was performed on the scaled data for the variable genes, and top principal components were chosen based on Seurat’s ElbowPlot (at least 15 PCs were used in all cases). Cells were clustered in PCA space using Seurat’s FindNeighbors on top principal components with default parameters, followed by FindClusters with resolution = 1.0. Variation in the cells was visualized by t-SNE (t-distributed stochastic neighbor embedding) on the top principal components.

For each dataset, upregulated genes in each cluster were identified using the VeniceMarker tool from the Signac package v0.0.7 from BioTuring (https://github.com/bioturing/signac). Cell types were manually assigned to each cluster by looking at the top most significant upregulated genes and comparing them to canonical markers used in literature to identify different cell types and to reference gene lists established in Uzquiano et al 2022. In a few cases, clusters were further subclustered to assign identities at higher resolution. Specifically:

Mito210 one month, cluster 18 was split into subcortical neurons and Cajal Retzius.

Mito210 three months batch 1, cluster 5 was split into CPN and unspecified PN, clusters 9 and 12 were each split into CFuPN and CPN, clusters 13 and 14 were each split into oRG and IP, and cluster 12 was split into CFuPN and CPN.

Mito210 three months batch 2, clusters 4, 9, and 10 were each split into CPN and CFuPN, and clusters 15 and 16 were each split into the oRG and IP.

PGP1 three months, clusters 13 and 15 were each split into oRG and IP, clusters 11 and 17 into CPN and CFuPN, and cluster 10 into PN and aRG

See metadata on Single Cell Portal (https://singlecell.broadinstitute.org/single_cell/study/SCP1964) to view cluster numbers.

To compare cell type proportions between control and mutant organoids, for each cell type present in a dataset, the glmer function from the R package lme4 v1.1-23 (Bates et al., Journal of Statistical Software, 2015) was used to estimate a mixed-effect logistic regression model. The output was a binary indicator of whether cells belong to this cell type, the control or mutant state of the cell was a fixed predictor, and the organoid that the cell belonged to was a random intercept. Another model was fit without the control-versus-mutant predictor, and the anova function from the stats core R package v3.6.3 was used to compare the two model fits. False discovery rate (FDR) was calculated with the Benjamini-Hochberg approach (Benjamini et. al., 1995).

To compare the proportion of proliferating oRG between conditions, Seurat’s CellCycleScoring function was used with default parameters, and any cells with an S score or G2M score above 0.5 was marked as proliferating. The significance of any difference in proportion was evaluated using a mixed-effect logistic regression model as above.

### Differential expression analysis

Differential genes between control and mutant organoids were assessed by subsetting each dataset into cells belonging to each cell type. Reads were then summed across cells of each type in each organoid (to create a pseudobulk per cell type). Genes with less than 10 total reads were excluded, and DESeq2 v1.26 (Love et al., Genome Biol, 2014) was used to calculate DEGs, with each organoid as a sample (Lun and Marioni, Biostatistics, 2017).

### Comparison of differentially expressed genes between control and mutant organoids to genes changing over time in organoid development

Organoid DEG signatures (above) were compared to genes changing in control organoids over time. To calculate the latter list, data from (Uzquiano et al., bioRxiv, 2022) were utilized. As in that paper, data from each time point (23 days, one month, 1.5 months, two months, three months, and four months) across all cell lines and organoids used in that paper, were corrected for ambient RNA expression using the decontX function from the Celda R package, version 1.6.1 (Yang et al., Genome Biol, 2020), with each organoid treated as its own sequencing batch. For each cell type, DESeq2 was used to identify genes that are differentially expressed across organoid age, in days. Two lists were made: one using data from 23 days, one month, and 1.5 months, to compare to the *PTEN* organoids at one month, and a second DEG list from the two months, three months, and four months timepoints, to compare to the *PTEN* organoids at 3 months.

DEG lists from control vs *PTEN* mutant organoids were compared to DEG lists across time using two methods. In both cases, DEG lists were ranked by degree of differentiation [-log10(p value) * sign(effect)] and then compared. First, they were compared using the improved rank-rank hypergeometric overlap test [RRHO2 R package v1.0; (Cahill et al., Sci Rep, 2018; Plaisier et al., Nucleic Acids Res, 2010)]. Second, they were compared using a Spearman correlation via the cor.test function from the stats core R package v3.6.3, and the rho statistic was reported.

### Slide-seqV2

For Slide-seqV2, Mito210 organoids at two months were embedded in OCT (Tissue Tek) and cryosectioned at 10 μm thickness. Sections were transferred to a custom-made array of densely packed barcoded beads (termed ‘pucks’) for Slide-seqV2 experiments. Library construction was performed as described (Stickels et al., Nat Biotechnol, 2021). Briefly, first-strand synthesis was performed by incubating the puck with tissue sections in the reversetranscription solution followed by tissue digestion, second-strand synthesis, library amplification, and purification. The Slide-seqV2 libraries were sequenced on a NovaSeq with an estimated ~100 million reads per puck. The data pertaining to control organoids have been reported previously in (Uzquiano et al., Cell, 2022).

### Slide-seq V2 data analysis

Sequencing reads were aligned to the GRCh38 human reference genome and analyzed by the Slide-seq pipeline (https://github.com/MacoskoLab/slideseq-tools) as previously described (Stickels et al., Nat Biotechnol, 2021). Since organoid sizes were much smaller than the Slide-seq pucks, images were manually cropped to the edges of the organoids and beads outside of the images were excluded.

Cell type decomposition was performed by Robust Cell Type Decomposition (RCTD; (Cable et al., Nat Biotechnol, 2022)). Briefly, cell type profiles learned from control organoid scRNA-seq data corresponding to the same time point and cell line (Uzquiano et al., Cell, 2022) were used to decompose mixtures from the Slide-seqV2 data. First, for each timepoint, a reference was made by using the age-matched scRNA-seq dataset. Next, the RTCD package fit a statistical model to estimate the mixture and cell type identities at each bead of the Slide-seqV2 data. We restricted our analysis to beads with more than 200 UMIs. Finally, gene set expression of the top 50 genes from each cell type signature as defined in (Uzquiano et al., Cell, 2022) was visualized by plotting 100x the sum of the reads for all genes in the gene set that appeared in the Slide-seqV2 data, normalized by the number of UMI for each bead.

### Bulk RNA-seq

For bulk RNA-seq, one month PGP1 *PTEN* organoids (n = 3 separate organoids for heterozygous mutant and isogenic control) were used. We extracted total RNA using Quick-RNA Mini Prep (Zymo Research) and followed the vendor protocol with a DNase treatment step. We constructed RNA-seq libraries using the Smart-seq2 protocol (Picelli et al., Nat Protoc, 2014), with minor modifications as follows: 1) we used 2 ng total RNA as input, 2) we used 0.1 ng cDNA as input, and made the NexteraXT (Illumina) libraries using half of the standard volume. We pooled libraries from different samples based on molar concentration, and sequenced them on a Nextseq500 instrument (Illumina) with 50 bases for read 1, 25 bases for read 2 and 8 bases each for Index 1 and 2. We performed the same experiment with Mito210 mutant and control organoids; however, the number of differentially-expressed genes between mutant and control was not sufficient to allow the RRHO2 analysis. Therefore, we did not proceed with the analysis of the Mito210 experiment.

For comparison of differentially expressed genes between control and mutant organoids to genes changing over time in organoid development, binary base call (BCL) files were converted to FASTQ files using the bcl2fastq (RRID:SCR_015058) package version 2.20.0. The reads were then aligned to human GRCh38 reference and the genes were quantified through the RSEM (Li and Dewey, BMC Bioinformatics, 2011) version 1.3.0 pipeline using the STAR (Dobin et al., Bioinformatics, 2013) version 2.7.10a aligner (using the following parameters: --star, --star-gzipped-read-file, --num-threads 4, –paired-end). DEGs between control and *PTEN* heterozygous mutant were identified using R (v4.0.3) package DESeq2 (v1.30.1) (Love et al., Genome Biol, 2014). Gene quantification results were loaded with tximport function (tximport package v1.18.0; (Soneson et al., F1000Res, 2015) and converted to DESeq data (DESeqDataSetFromTximport function from DESeq2). A negative binomial Wald test was applied to find DEGs between the two conditions (DESeq function with the default settings). To compare whether the obtained DEGs show concordance with accelerated or delayed differentiation, we identified DEGs that vary in expression across time, using all cells in a pseudobulk, and used RRHO2 to compare DEG lists, as in the above section “Comparison of differentially expressed genes between control and mutant organoids to genes changing over time in organoid development”.

### Cell lysis and filter-aided sample preparation (FASP) digestion for mass spectrometry

For one month (35 *d.i.v*.) *and* two months (70 *d.i.v.*) Mito210 *PTEN* organoids, four heterozygous mutant and four control and three heterozygous mutant and three control organoids were used, respectively. Cells were placed into microTUBE-15 (Covaris) microtubes with TPP buffer (Covaris) and lysed using a Covaris S220 Focused-ultrasonicator instrument with 125 W power over 180 seconds, at 10% max peak power. Upon precipitation with chloroform/methanol, extracted proteins were weighed and digested according to the FASP protocol (100 μg). Briefly, the 10K filter was washed with 100 μl of triethylammonium bicarbonate (TEAB). Each sample was added, centrifuged, and the supernatant discarded. Then, 100 μl of Tris(2-carboxyethyl) phosphine (TCEP) 37 was added for one hour, centrifuged, and the supernatant discarded. While shielding from light, 100 μl IAcNH2 was added for 1 hour followed by centrifugation, and the supernatant was discarded. Next, 150 μl 50 mM TEAB + Sequencing Grade Trypsin (Promega) was added and left overnight at 38° C, upon which the samples were centrifuged and supernatant collected. Lastly, 50 μl 50 mM TEAB was added to the samples, followed by spinning and supernatant collection. The samples were then transferred to HPLC.

### TMT Mass tagging protocol peptide labeling

The TMT (Tandem Mass Tag) Label Reagents were equilibrated to RT and resuspended in anhydrous acetonitrile or ethanol (for the 0.8 mg vials 41 μl were added, for the 5 mg vials 256 μl were added). The reagent was dissolved for 5 minutes with occasional vortexing. TMT Label Reagent (41 μl, equivalent to 0.8 mg) was added to each 100-150 μg sample. The reaction was incubated for one hour at RT. Reaction was quenched by adding 8 μl of 5% hydroxylamine to the sample and incubating for 15 minutes. Samples were combined, dried in a SpeedVac (Eppendorf), and stored at −80° C.

### Hi-pH separation and mass spectrometry analysis

Before submission to LC-MS/MS (Liquid Chromatography with tandem mass spectrometry), each experiment sample was separated on a Hi-pH column (Thermo Fisher Scientific) according to the vendor’s instructions. After separation into 40 fractions, each fraction was submitted for a single LC-MS/MS experiment, performed on a Lumos Tribrid (Thermo Fisher Scientific) equipped with 3000 Ultima Dual nanoHPLC pump (Thermo Fisher Scientific). Peptides were separated onto a 150 μm inner-diameter microcapillary trapping column, packed first with approximately 3 cm of C18 Reprosil resin (5 μm, 100 Å, Dr. Maisch GmbH) followed by PharmaFluidics micropack analytical column 50 cm. Separation was achieved through applying a gradient from 5-27% ACN in 0.1% formic acid over 90 minutes at 200 nl per minute. Electrospray ionization was enabled through applying a voltage of 1.8 kV using a home-made electrode junction at the end of the microcapillary column and sprayed from stainless-steel tips (PepSep). The Lumos Orbitrap was operated in data-dependent mode for the MS methods. The MS survey scan was performed in the Orbitrap in the range of 400 – 1,800 m/z at a resolution of 6 × 104, followed by the selection of the 20 most intense ions (TOP20) for CID-MS2 fragmentation in the Ion trap using a precursor isolation width window of 2 m/z, AGC setting of 10,000, and a maximum ion accumulation of 50 ms. Singly-charged ion species were not subjected to CID fragmentation. Normalized collision energy was set to 35 V and an activation time of 10 ms. Ions in a 10 ppm m/z window around ions selected for MS2 were excluded from further selection for fragmentation for 90 seconds. The same TOP20 ions were subjected to HCD MS2 event in Orbitrap part of the instrument. The fragment ion isolation width was set to 0.8 m/z, AGC was set to 50,000, the maximum ion time was 150 ms, normalized collision energy was set to 34 V and an activation time of 1 ms for each HCD MS2 scan.

### Mass spectrometry data generation

Raw data were submitted for analysis in Proteome Discoverer 2.4 (Thermo Fisher Scientific) software. Assignment of MS/MS spectra was performed using the Sequest HT algorithm by searching the data against a protein sequence database including all entries from the Human Uniprot database (Bairoch and Apweiler, Nucleic Acids Res, 1999; UniProt Consortium, Nucleic Acids Res, 2018) (Bairoch & Apweiler, 1999, Consortium, 2018) and other known contaminants such as human keratins and common lab contaminants. Sequest HT searches were performed using a 10 ppm precursor ion tolerance and requiring each peptide’s N-/C termini to adhere with Trypsin protease specificity, while allowing up to two missed cleavages. 16-plex TMTpro tags on peptide N termini and lysine residues (+304.207 Da) was set as static modifications while methionine oxidation (+15.99492 Da) was set as variable modification. A MS2 spectra assignment false discovery rate (FDR) of 1% on protein level was achieved by applying the target-decoy database search. Filtering was performed using Percolator (64 bit version) (Käll et al., J Proteome Res, 2008). For quantification, a 0.02 m/z window centered on the theoretical m/z value of each of the 6 reporter ions and the intensity of the signal closest to the theoretical m/z value was recorded. Reporter ion intensities were exported in the result file of Proteome Discoverer 2.4 search engine as Excel tables. The total signal intensity across all peptides quantified was summed for each TMT channel, and all intensity values were normalized to account for potentially uneven TMT labeling and/or sample handling variance for each labeled channel.

### Mass spectrometry data analysis

Potential contaminants were filtered out and proteins supported by at least two unique peptides were used for further analysis. We kept proteins that were missing in at most one sample per condition. Data were transformed and normalized using variance stabilizing normalization using the DEP package of Bioconductor (Zhang et al., Nat Protoc, 2018). To perform statistical analysis, data were imputed for missing values using random draws from a Gaussian distribution with width 0.3 and a mean that is down-shifted from the sample mean by 1.8. To detect statistically significant differential protein abundance between conditions, we performed a moderated t-test using the LIMMA package of Bioconductor (Ritchie et al., Nucleic Acids Res, 2015), employing an FDR threshold of 0.1. GSEA was performed using the GSEA software (Subramanian et al., Proc Natl Acad Sci U S A, 2005). GO and KEGG pathway annotation were utilized to perform functional annotation of the significantly regulated proteins. GO terms and KEGG pathways with FDR q-values < 0.05 were considered statistically significant. Correlation between mutant effect (e.g., *PTEN*^+/−^ *vs* Control at 1 month) and time effect (e.g., 2 *vs* 1 month in Control) was calculated using Pearson correlation. For *PTEN* 2 vs 1 month, changes in protein levels in heterozygous and control organoids were compared to one another with a signed paired Wilcoxin rank test, using stat_compare_means from the ggpubr R package (https://rpkgs.datanovia.com/ggpubr/).

### Calcium Imaging

Organoids were transduced with pAAV-CAG-SomaGCaMP6f2 (Addgene, #158757) by pipetting 0.2 μl of stock virus to 500 μl of Cortical Differentiation Medium IV (CDM IV, without matrigel) in one well of a 24 well dish containing a single organoid. On the next day, each organoid was transferred to a 6-well plate filled with 2 ml of fresh medium. On the third day after transduction, organoids were transferred to low-attachment 10-cm plates, and on the seventh day, medium was switched to BrainPhys (5790 STEMCELL Technologies) supplemented with 1% N2 (17502-048 Thermo Fisher), 1% B27 (17504044 Thermo Fisher), GDNF (20 ng/ml, Cat. No. 78139 STEMCELL Technologies), BDNF (20 ng/ml, 450-02 Peprotech), cAMP (1mM, 100-0244 Stemcell Technologies), Ascorbic acid (200 nM, Cat. No. 72132 STEMCELL Technologies), and laminin (1 μg/ml, 23017015 Life Technologies). Organoids were cultured in BrainPhys for at least 2 weeks before imaging.

Brain organoids were randomly selected and transferred to a recording chamber containing BrainPhys. Imaging was performed using a confocal scanner (CSU-W1, Andor confocal unit attached on an inverted microscope [Ti-Eclipse, Nikon]), while the organoids were kept at 37°C using a heating platform and a controller (TC-324C, Warner Instruments). We used a 10x objective (Plan Apo λ, 10x/0.45), resulting in a field of view of 1.3 x 1.3 mm^2^ and a pixel size of 0.6 μm. Imaging took place in fast-time-lapse mode, with an exposure time of 100ms, resulting in an acquisition rate of approximately 8 frames/sec. Spontaneous activity was recorded in three different z-planes, for at least 22 minutes of baseline activity in total (with no pharmacology treatment).

Stock solutions of 2,3-dioxo-6-nitro-1,2,3,4-tetrahydrobenzo[*f*]quinoxaline-7-sulfonamide disodium salt (NBQX disodium salt, Abcam; 100 mM) and Tetrodotoxin citrate (TTX, Abcam; 10 mM) were prepared in ddH_2_O. Bath application of NBQX (antagonist of AMPA/kainate glutamate receptors) and TTX (voltage-gated sodium-channel antagonist) was applied to achieve a final bath concentration of 20 μM and 2 μM, respectively.

Data were converted from nd2 format to tiff, and automated motion correction and cell segmentation were performed using Suite2p (Pachitariu et al., bioRxiv, 2017), followed by manual curation of segmented cells (we examined the spatial footprint and temporal characteristics of each candidate cell, as well as manually adding neurons with clear cellbody morphology; see Fig. 2d). Then, mean raw fluorescence for each cell was measured as a function of time.

### Analysis of calcium imaging data

Analysis was done using in-house MATLAB scripts. Raw calcium signals for each cell, F(t), were converted to represent changes from baseline level, ΔF/F(t), defined as (*F*(t) – *F*o(t))/*F*o(t). The time-varying baseline fluorescence, *F*o(t), for each cell was a smoothed fluorescence trace obtained after applying a 10-second-order median filter centered at t. Calcium events elicited by action potentials were detected based on a threshold value given by their peak amplitude (5 times the standard deviation of the noise value) and their first time derivative (2.5 times the standard deviation of the noise value).

The analysis of network bursting was performed based on the population-averaged calcium signal along all segmented cells. A peak in the population signal was considered a network burst if it met the following criteria: (1) the peak amplitude was greater than 10 times the standard deviation of the noise value, (2) a set of bursting cells composed of at least 20% of total cells were active during that population calcium transient, and (3) a cell was considered part of the set of bursting cells only if it participated in at least 50% of the network bursts.

The peaks of the network bursts were used to measure the inter-burst interval (IBI), and the frequency was obtained from the average IBI. The burst half-width was also measured from the population-averaged calcium signal by calculating the width of the transient at 50% of the burst peak amplitude.

## Notes

https://singlecell.broadinstitute.org/single_cell/study/SCP1964

